# Single-cell Spatial Metabolic and Immune Phenotyping of Head and Neck Cancer Tissues Identifies Tissue Signatures of Response and Resistance to Immunotherapy

**DOI:** 10.1101/2023.05.30.540859

**Authors:** Niyati Jhaveri, Bassem Ben Cheikh, Nadezhda Nikulina, Ning Ma, Dmytro Klymyshyn, James DeRosa, Ritu Mihani, Aditya Pratapa, Yasmin Kassim, Sidharth Bommakanti, Olive Shang, Yan He, Yi Zheng, James Monkman, Caroline Cooper, Ken O’Byrne, Bhaskar Anand, Michael Prater, Subham Basu, Brett G.M. Hughes, Arutha Kulasinghe, Oliver Braubach

## Abstract

Head and neck squamous cell carcinomas (HNSCC) are the seventh most common cancer and represent a global health burden. Immune checkpoint inhibitors (ICIs) have shown promise in treating recurrent/metastatic cases, with durable benefit in ∼30% of patients. Current biomarkers for head and neck tumors are limited in their dynamic ability to capture tumor microenvironment (TME) features, with an increasing need for deeper tissue characterization. Therefore, new biomarkers are needed to accurately stratify patients and predict responses to therapy. Here, we have optimized and applied an ultra-high plex, single-cell spatial protein analysis in HNSCC. Tissues were simultaneously analyzed with a panel of 101 antibodies that targeted biomarkers related to tumor immune, metabolic and stress microenvironments. Our data uncovered a high degree of intra-tumoral heterogeneity intrinsic to head and neck tumors and provided unique insights into the biology of the tumor. In particular, a cellular neighborhood analysis revealed the presence of 6 unique spatial tumor-immune neighborhoods enriched in functionally specialized immune cell subsets across the patient tissue. Additionally, functional phenotyping based on key metabolic and stress markers identified four distinct tumor regions with differential protein signatures. One tumor region was marked by infiltration of CD8+ cytotoxic T cells and overexpression of BAK, a proapoptotic regulator, suggesting strong immune activation and stress. Another adjacent region within the same tumor had high expression of G6PD and MMP9, known drivers of tumor resistance and invasion respectively. This dichotomy of immune activation-induced death and tumor progression in the same sample demonstrates the heterogenous niches and competing microenvironments that underpin clinical responses of therapeutic resistance. Our data integrate single-cell ultra-high plex spatial information with the functional state of the tumor microenvironment to provide insights into a partial response to immune checkpoint inhibitor therapy in HNSCC. We believe that the approach outlined in this study will pave the way towards a new understanding of TME features associated with response and sensitivity to ICI therapies.

## INTRODUCTION

Head and neck squamous cell carcinomas (HNSCC) are tumors that develop in the lip, oral cavity, larynx, salivary glands, nose, sinuses, or the skin of the face. They represent the seventh most common cancer globally with >650,000 new cases and >300,000 deaths annually (1). The primary risk factors include tobacco use, excessive consumption of alcohol, and infection with human papilloma virus (HPV) (2). The current treatment regimen for HNSCC remains surgery and radiotherapy with/without chemotherapy, but resistance and tumor recurrence contribute to an overall poor prognosis (3). While immunotherapies have been approved in the recurrent/metastatic setting for HNSCC, and some have shown to provide a durable benefit, better predictive biomarkers are needed to identify those HNSCC patients that will achieve therapeutic benefits (4–6).

Immunosuppression is a characteristic hallmark of HNSCC and analysis of immune checkpoints PD-1/PD-L1, FOXP3+ regulatory T cells and CD8+ tumor infiltrating lymphocytes in the tumor microenvironment has been explored extensively (4). However, there are no validated biomarkers to predict immunotherapy responsiveness across HNSCC patients, likely due to the high degree of intra and inter-tumoral heterogeneity. A promising avenue in this regard is the recent emergence of spatial biomarkers that seem to better predict patient outcomes to immunotherapy. For example, a retrospective analysis of immune cell neighborhood frequencies in cutaneous T cell lymphomas have highlighted the potential for spatial analysis of tumor-associated immune cells as a means to stratify immunotherapy-sensitive and resistant cohorts (7). Several additional studies now support the utility of spatial biomarkers (8–11).

To date, most studies on spatial biomarkers have been focused on immune cell populations within the tumor microenvironment. However, tumor biology is much more complex and spans a wide range of biological mechanisms and pathways (12,13). For example, cellular metabolism is critical to the functioning of *all* cells and metabolic reprogramming in cancer plays an important role in tumor progression, immune activation, and metastasis (14–17). Studies investigating metabolic tissue signatures in HNSCC are limited and lack spatial context that would be critical to analyze the metabolome at the single-cell level (18). Coupling spatial analyses of the tumor immune microenvironment with information about cellular metabolism and stress could thus provide valuable new insight into the HNSCC biomarker landscape and identify predictive signatures of response and resistance.

In the present study, we developed an ultra-high plex single-cell spatial protein analysis of HNSCC. Our workflow (**Supplementary Figure 1A**) is based on CO-Detection by indEXing (CODEX) (19,20). CODEX employs cocktails of DNA-tagged antibodies to stain tissue, which is then followed by iterative steps of hybridization with complementary, fluorescently labeled probes for imaging. This results in the generation of imaging data with dozens of antibody readouts, thereby enabling the extraction of high-parameter, spatially resolved single-cell data from solid tissue samples (21–23). Crucially, CODEX is based on optical imaging and thus offers established technical advantages of high resolution, scalability of imaging area, as well as wide-ranging flexibility. The commercialized version of CODEX, the PhenoCycler^®^ (Akoya Biosciences, The Spatial Biology Company, Marlborough, MA, USA) is widely deployed and the workflow is seamlessly compatible with a wide range of other applications. In this study, we applied a 101-plex antibody panel spanning biomarkers from the hallmarks of cancer (12,13), including immune checkpoints, tumor-promoting inflammatory biomarkers, markers related to angiogenesis, invasion, and metastasis, as well as pathways implicated in the deregulation of cellular energetics, proliferation, evading growth suppression and apoptosis. The sum of our data allows us to provide a comprehensive account of not only the tumor immune microenvironment, but also provide insight into the metabolic state and biology of different regions within the tumor; in doing so, our data describe an intricate interplay between immune infiltration of certain tumor regions versus the metabolic and stress responses of tumor cells within these regions.

## MATERIALS AND METHODS

### Tissues and Experimental Procedures

#### Panel Development

Formalin-fixed paraffin-embedded human tonsil tissue was purchased from BioIVT (Westbury, NY) for development and validation of the 101-plex panel.

#### Tumor Phenotyping

Retrospective HNSCC tissues were collected before treatment and stored as FFPE blocks from the Royal Brisbane and Women’s Hospital (RBWH) with Human Research Ethics HREC Approval No. LNR/2020/QRBW/66744 and The University of Queensland ratification. Patients were treated with pembrolizumab or nivolumab and categorized based on response to therapy according to RECIST 1.1 including complete response (CR), partial response (PR), stable disease (SD), and progressive disease (PD). Serial sections of hematoxylin and eosin (H&E) staining were prepared by Pathology Queensland and reviewed by a pathologist for tumor/stroma demarcation. A HNSCC tissue from a patient with a partial response was analyzed by spatial phenotyping.

#### Sectioning

Following equilibration of blocks at room temperature, 5µm thick sections were cut using a Leica RM2125 RTS microtome and sectioned onto Poly-L-lysine coated slides (Leica Surgipath Apex Superior adhesive slides, #3800081). Slides were dried at room temperature overnight and stored at 4°C for long-term storage.

#### Reagents

Commercially available oligo-conjugated antibodies were obtained from Akoya Biosciences (Marlborough, MA). Please refer to **Supplementary Table 1** for antibody and catalog information. Additional antibodies were obtained from commercial vendors as indicated and conjugated to custom oligo barcodes according to instructions in Akoya’s antibody conjugation kit (Conjugation kit, Akoya, #7000009). Briefly, antibodies were reduced to expose thiol groups in the hinge region followed by conjugation to unique oligonucleotides. After multiple rounds of purification, oligo-conjugated antibodies were eluted and stored in Storage Buffer at 4°C. Complementary oligo-conjugated fluorophore reporters were obtained from Akoya Biosciences.

#### Sample Processing

Slides were baked for 1 hour at 55°C on a slide warmer (Premiere XH-2004) and dewaxed through two successive rounds of incubation in Histochoice Clearing Agent (VWR, PN# H103-4L) for 5 minutes each. Subsequently, the FFPE sections were dehydrated by sequentially incubating in 100% ethanol (Sigma Aldrich, PN# 79317-16GA-PB), 90% ethanol, 70% ethanol, 50% ethanol and 30% ethanol twice each for 5 minutes. Slides were then rinsed three times in distilled water for 5 minutes to ensure no carryover of ethanol before target retrieval. Heat-mediated epitope retrieval was performed by incubating slides in a beaker containing Tris EDTA solution (pH 9; Dako, #S2367) in a pressure cooker for 20 minutes. After 20 minutes, the beaker was removed from the pressure cooker and left to equilibrate at room temperature for 30 minutes. Slides were then rinsed through two rounds of incubation in distilled water for 2 minutes and stored in Hydration Buffer from the PhenoCycler Staining Kit (Akoya Biosciences, MA, #7000008) until ready for staining.

For antibody staining, an antibody cocktail was prepared with optimal dilutions of each antibody in a buffer containing N, J, S and G blockers (Akoya Biosciences, MA, #7000008). Samples were first allowed to equilibrate to room temperature in Staining Buffer for 20-30 minutes, followed by incubation in a pre-blocking solution made of N, J and S blockers in Staining Buffer. The first antibody cocktail (containing the first 50 antibodies) was subsequently added to the slide and slides were incubated overnight at 4°C. On the next day, slides were rinsed with Staining buffer and fixed with 1.6% PFA (Electron Microscopy Sciences, PN# 15710) in Storage buffer solution. A second round of antibody staining (with the remaining antibodies) was performed with incubation at room temperature for 3 hours. After washing with Staining buffer and fixation of the second set of antibodies in 1.6 % PFA, samples were fixed in ice cold methanol for 5 minutes at 4°C. A third round of fixation with PhenoCycler Fixative reagent was applied and slides were stored in Storage Buffer at 4°C until ready to image.

#### Reporter Plate Preparation and Imaging

Reporter Stock solution was prepared as described in the PhenoImager Fusion User Guide (Version 1.0.6_RevI) with 10X Buffer (#7000001), Assay Reagent (#7000002) and Nuclear Stain (#7000003) from Akoya Biosciences. Individual tubes of 3 reporters/cycle, diluted in reporter stock solution, were prepared based on the cycle information in **Supplementary Table 2**. Subsequently, a 96 well plate was prepared with 1 well/cycle containing the corresponding working reporter solution for that cycle. Blank cycles containing only reporter stock solution without any added reporters were included as the first and last cycle in each run for subtraction of autofluorescence background.

Antibody-stained slides were next equilibrated at room temperature in 1X Buffer for at least 10 minutes. The flow cell (Akoya Biosciences, #240204) was assembled onto the slide as described in the user guide. The slides were incubated in 1X Buffer again for 10 minutes to ensure secure sealing of the flow cell to the slide. The slide was then transferred to the flow cell carrier and imaged on a PhenoCycler-Fusion (Akoya Biosciences, MA) using the following exposure settings: DAPI – 1 ms, ATTO550 channel – 150 ms, AF647 channel – 150 ms and AF750 channel – 150 ms. Integration of the PhenoCycler automated fluidics cycler with the PhenoImager Fusion imaging system automated the entire process of reporter hybridization, imaging and dehybridization to capture whole slide images of three markers (+DAPI for nuclear staining) at a time. After images of all cycles had been acquired, the final QPTIFF file containing a composite image of all markers was viewed using the QuPath software (https://qupath.github.io/), where each channel can be turned on and off individually or collectively to reveal the spatial expression pattern of the marker(s) of interest.

### Bioinformatic Analysis

Computational image analysis was used to identify different cell types and characterize their spatial distribution in the tissue. The image analysis workflow is described in **Supplementary Figure 1B**. Before initiating data analysis, quality control was performed on each individual marker image by visual assessment and filtering was conducted to exclude tissue regions with poor staining outcomes. The subsequent first step of the image analysis then consisted of cell segmentation, which identified individual cells across the tissue by determining their location and surface. The second step consisted of phenotyping the segmented cells based on their combinatorial antibody labeling intensities. The final step involved characterizing the spatial organization of the different cell types and their spatial interactions in local cellular neighborhoods.

#### Segmentation

Nuclear segmentation was first performed using the deep learning method StarDist (24) applied to the DAPI image of the tissues. Contrast Limited Adaptative Histogram Equalization (CLAHE) was applied as a preprocessing step to enhance image contrast during training and testing. Segmentation of the cytoplasm was then determined from nuclear expansion by morphological dilation applied to the labelled nuclear mask, and the centroid of each cell was defined by the x-y coordinates of the nuclear object centroid in the image.

#### Phenotyping

After cell segmentation, the average staining intensity for each protein was calculated for each individual cell from the corresponding expression compartment. For proteins localized to cell nuclei (e.g. Ki67, PCNA, FOXP3), the average intensity was calculated from the nuclear mask obtained with StarDist segmentation. If the protein was localized to the cell membrane (e.g. CD4, CD3, E-cadherin), the average intensity was calculated from the membrane mask. This step produced an expression table where cells are listed with their protein expressions and locations in the tissue. Each protein expression was then z-scored across all cells, such that every biomarker’s expression mean equaled 0 with a standard deviation equal to 1. For cell phenotyping, a particular subset of proteins and cells was selected depending on the question asked in our study, e.g. immune-cell subtyping or metabolic phenotyping.

Unsupervised clustering was performed using the Leiden algorithm implementation from the Scanpy toolkit (25) and the GPU-accelerated Rapids package (26). Leiden *resolutions* equal to 1-6 were tested and the optimal *resolution* of 4 was chosen for manual cluster annotation of cell phenotypes represented in hierarchical clustering heatmaps. Clusters with similar expression profiles were combined into one phenotype and a new heatmap with re-averaged protein expressions and cell phenotypes was generated (**Figures 2A, 3C and 5A**).

**Figure 1:**
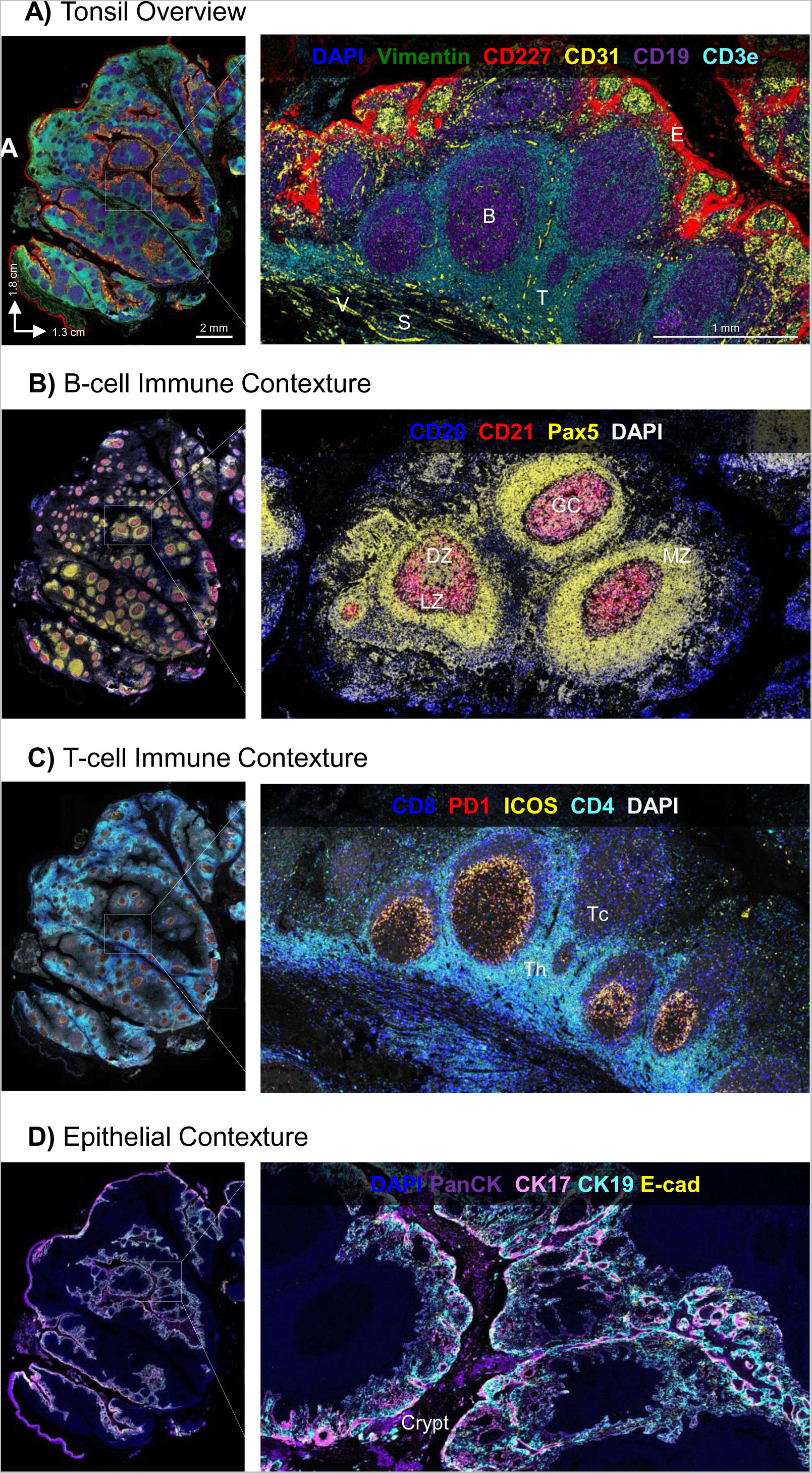
Whole-slide imaging of key cell phenotypic markers at single cell resolution reveals the complex topography of human tonsil tissue. A) An overview of human FFPE tonsil tissue stained with Vimentin (green), CD227/MUC1 (red), CD31 (yellow), CD19 (purple) and CD3e (cyan) marks stroma (S), epithelia (E), vasculature (V), B cells (B) and T cells (T), respectively, within this tissue. B) Zooming into germinal centers (GCs) with imaging of CD20 (blue), CD21 (red) and Pax5 (yellow), reveals the spatial organization of B cells with dark zones (DZs), light zones (LZs) and mantle zones (MZs) comprising distinct lymphoid follicles. C) Multiplexed imaging of T cell markers CD8 (blue), PD1 (red), ICOS (yellow) and CD4 (cyan) further unveils the distribution and localization of helper T cells (Th) and cytotoxic T cells (Tc) across the tissue. D) Stratified squamous epithelium identified by positive staining for PanCK (purple), CK17 (pink), CK19 (cyan) and E-cadherin (yellow) invaginates to form tonsillar crypts in tonsil.

**Figure 2:**
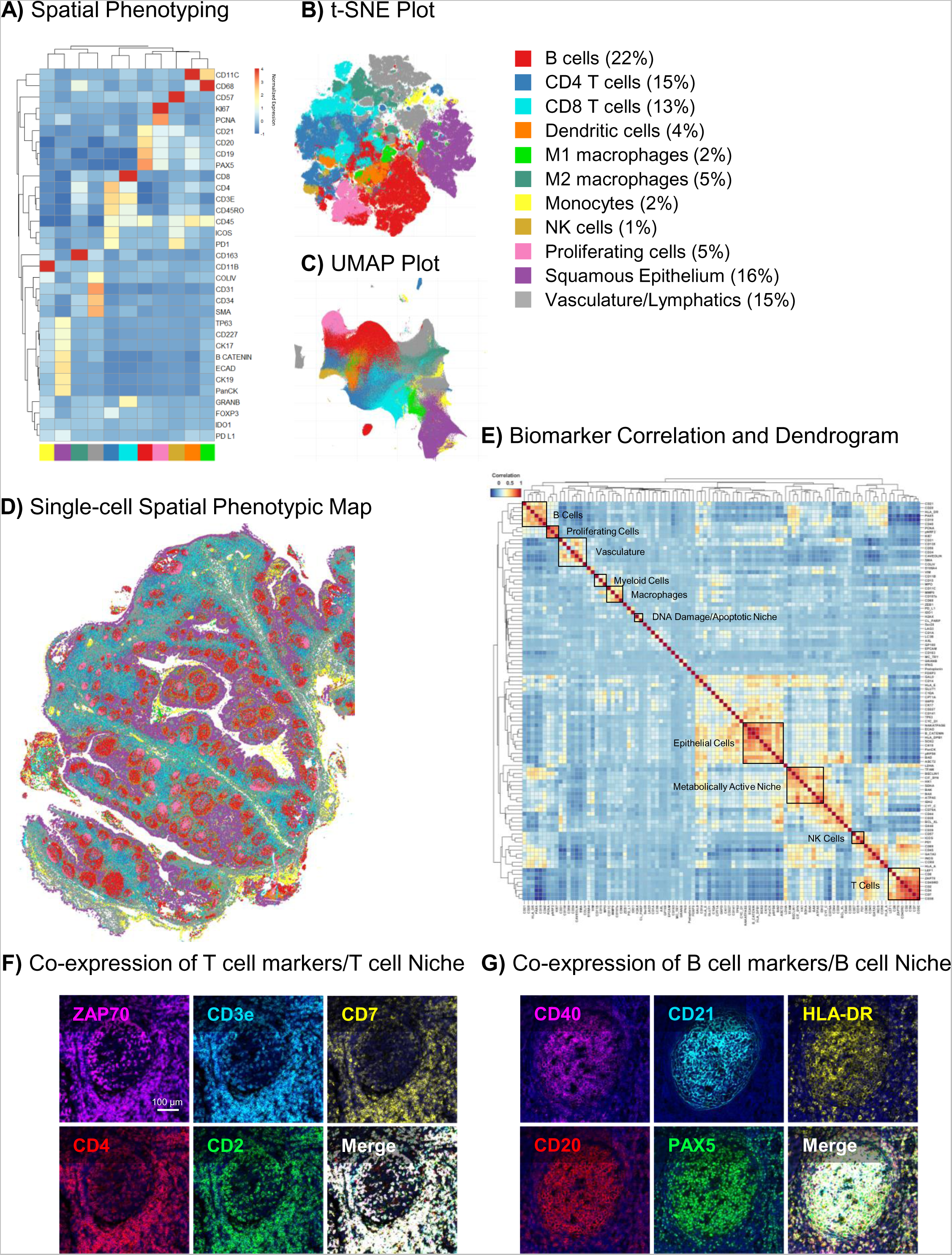
Single-cell spatial phenotyping of a human FFPE tonsil tissue reveals 11 distinct cell types. A) The heatmap (with markers listed on the right) shows a curated clustering dendrogram with cell types (columns) corresponding to the legend as indicated on the far right. B) and C) show, respectively, the t-SNE and UMAP plots of all 2,253,539 cells in a human tonsil colored by phenotype. D) A single-cell spatial map shows the distribution of the 11 distinct cell phenotypes across the entire tissue and reveals the organization of germinal centers, T cell zones and tonsillar crypts. E) A marker correlation heatmap indicates cell lineages and functional niches with varying metabolic, immune, and stress signatures. F) Co-expression of T cell markers as seen in the marker correlation heatmap is validated in multiplex images of ZAP70, CD3e, CD7, CD4 and CD2 in tonsil tissue. G) Similarly, co-expression of B cells markers – CD40, CD21, HLA-DR, CD20 and PAX5 is shown in a germinal center, thus verifying the specificity of the panel.

**Figure 3:**
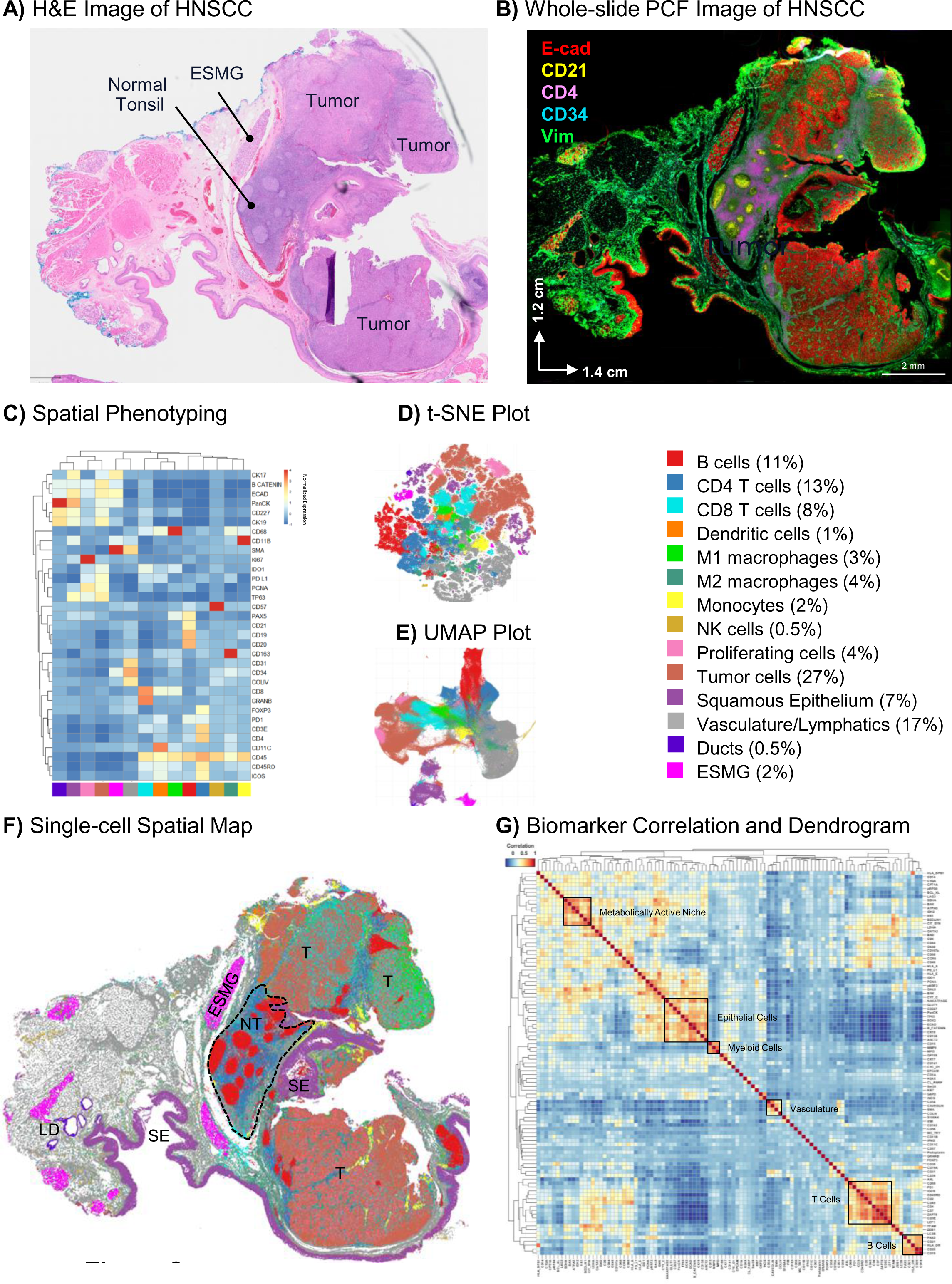
Interrogating single-cell spatial proteomics from a 101-plex phenotyping panel across a human FFPE head and neck squamous cell carcinoma. A) H&E-stained image of HNSCC showing pathologist’s annotations for normal tonsil region, tumor regions and esophageal submucosal glands (ESMG). B) Whole-slide imaging at single-cell resolution of epithelia (E-cadherin, red), B cells (CD21, yellow), T cells (CD4, pink), blood vessels (CD34, cyan) and stroma (Vimentin, green). C) A heatmap shows spatial phenotyping of the HNSCC into 14 distinct cell types (legend on far right). D) and E) show, respectively, the t-SNE and UMAP plots of all 856,676 cells in the tissue colored by phenotype. F) A spatial map illustrates the organization of the 14 cell phenotypes across the entire tissue and reveals a normal tonsil region (NT, dotted outline) with intact germinal centers and dense tumor regions (T). Esophageal submucosal glands (ESMG), lymphatic ducts (LD) and normal squamous epithelium (SE, purple) are also indicated. G) A marker correlation heatmap of all biomarkers indicates clustering and co-expression of certain markers.

Dimensionality reduction was applied for data visualization by means of UMAP and tSNE using Scanpy and Rapids implementations (25,26). UMAP parameters were set to 20 for the *number of neighbors*, 0.05 for *minimum distance*, and 0.5 for *spread* while tSNE parameters were set to 30 for *perplexity*, 12 for *early exaggeration*, and 200 for the *learning rate*. Furthermore, for the HNSCC tissue, regions of interest (ROIs) were annotated as Tumor 1, Tumor 2, Tumor 3, and Tumor 4, based on metabolic phenotyping obtained from the unsupervised clustering of key proteins involved in cellular metabolism and stress pathways. The number of cell types in each annotated region was then calculated, and the relative cell percentages were compared between the different tumor regions.

#### Cellular Neighborhood Analysis

The spatial interactions between different cell phenotypes were quantified using the Cellular Neighborhood method described previously (8). This method provides critical information about the relative abundance of cell types within the local neighborhood of a cell. In this study, the neighborhood of a cell was first defined using a Delaunay graph, where a cell N is considered to be a neighbor of cell A if they are connected in terms of Delaunay triangulation (see **Supplementary Figure 2A**). Then, for each cell, the cell-type percentages in its neighborhood were calculated. This step produces a neighborhood matrix where cells are characterized by the percentages of cell types in their local neighborhood. K-means clustering was then performed on all cells based on these percentages with k = {3,..,20}. The optimal number of cellular neighborhoods was determined as 6 based on Calinski-Harabasz criterion (27) (see **Supplementary Figure 2B**). The average neighborhood percentages were also calculated for each cellular neighborhood and plotted on a heatmap (see **Supplementary Figure 2C**).

## RESULTS

### Development of an ultra-high plex antibody panel for comprehensive spatial phenotyping of human FFPE tissues

High-plex single-cell spatial phenotyping of human tissues has led to valuable insights into the immune contexture that plays a critical role in tumor surveillance, regression, and recurrence (7,8). To date, these studies have mainly focused on the tumor immune microenvironment with antibody panels designed to identify immune cell lineages, immune activation states, immune checkpoints, as well as some tumor specific markers. The sheer complexity of tumor biology warrants a more comprehensive and multi-dimensional spatial phenotyping approach to analyze the tumor microenvironment. To this end, we developed an ultra-high plex antibody panel that encompasses antibodies for detection of not only immune and cancer cells, but also markers that identify cellular metabolism, apoptosis and stress, tumor invasion and metastasis as well as cellular proliferation and deregulation. We first evaluated the performance of this panel on human FFPE tonsil tissues.

#### Evaluating antibody panel performance on a tonsil tissue

All antibodies that are commercially available from Akoya Biosciences (**Supplementary Table 1**) have been previously validated in comparisons with orthogonal standard IHC staining in matching human FFPE tissues (19) (**Supplementary Figure 3a**); antibodies that were custom conjugated for this experiment (**Supplementary Table 1**) were qualitatively confirmed by means of comparing staining patterns with available datasets from the antibody suppliers. All antibodies were assessed for appropriate staining intensity and signal to noise ratios (**Supplementary Figure 3b**).

All antibodies were then assembled into an ultra-high plex panel of 101 markers and deployed on a human FFPE tonsil tissue for comprehensive spatial phenotyping and characterization of the tissue microenvironment (**Supplementary Figure 4**). Three markers were imaged per cycle with a total of 38 cycles of reporter hybridization, imaging, and de-hybridization to reveal the spatial localization of all 101 proteins across the entire tissue section. A representative image (**Figure 1a**) showing epithelia (CD227/MUC1), T cells (CD3e), B cells (CD19), stroma (Vimentin), and blood vessels (CD31) recapitulates regional and cellular tonsillar features such as germinal centers and tonsillar crypts.

A deeper dive into the spatial biology of the tonsil reveals additional distinct cellular subsets within the tissue microenvironment. First, we examined B cells (CD20+ CD21+ Pax5+ cells), which are responsible for production of antigen-specific immunoglobulins and the humoral arm of the adaptive immune response. Within secondary lymphoid tissues like tonsil, B lymphocytes aggregate to form distinct lymphoid follicles, which on activation expand to form prominent pale regions called germinal centers. In response to antigen exposure, rapidly proliferating B cells push primary follicle cells to the periphery, generating a mantle zone around the germinal center. The characteristic microenvironment of the germinal center is visible here in **Figure 1b** with densely packed B cells forming the dark zone and a sparsely populated region of B cells forming the light zone along with T follicular helper cells and follicular dendritic cells (28).

T lymphocytes are another critical component of adaptive immunity and are found in T cell rich zones bordering germinal centers where they play an important role in B cell selection and survival. Different subsets of T cells such as CD4+ follicular helper T cells, CD8+ cytotoxic T cells and FOXP3+ regulatory T cells have unique spatial expression patterns within tonsil in concordance with their functional significance. The complexity of heterogeneous T cell subsets and their regulation by immune checkpoints such as ICOS and PD-1 requires multiplexed analysis of all the relevant markers at single cell resolution as shown in **Figure 1c**.

Apart from immune subsets that comprise the majority of tonsil tissue, the luminal surface is lined by non-keratinizing stratified squamous epithelium (29) (**Figure 1d**). The epithelium (marked by cytokeratins and E-cadherin) invaginates and forms tonsillar crypts in response to antigenic stimulation. The structure of the crypt lined by alternating patches of reticulated epithelium mixed with lymphoid cells and patches of squamous epithelium facilitates the initial step of an adaptive immune response; the number of crypts correlate with the extent and reactivity of the lymphoid component.

An interactive dataset (https://dataset.akoyabio.com/PCF-Tonsil100plex/) shows additional layers of information that can be obtained from this tonsil data. Collectively, these data confirm the accuracy of our 101-plex panel to capture regional and cellular morphological features of human FFPE tonsil tissue. Moreover, they confirm the possibility of conducting deep characterization of cellular subsets *in situ* within single specimens. Combining the acuity of histology with high-parameter single-cell biomarker readouts provides critical insights into the structural and functional complexity of human tissues and the processes that maintain the balance between homeostasis and disease. We next proceeded to systematically analyze these data using an unsupervised bioinformatic single-cell analysis approach.

#### Unsupervised clustering and regional analysis of ultra-high plex staining data

The single-cell spatial organization of the tonsil tissue was evaluated by conducting an unsupervised analysis of a subset of biomarkers related to the immune system and tissue architecture. To this end, we performed nuclear segmentation with Stardist and unsupervised Leiden clustering of all 2.25 million cells present in the tonsil tissue (see methods for more details). Thirty-three biomarkers were included in this analysis, including cell lineage markers, immune checkpoints, and structural markers. As shown in the clustering dendrogram (**Figure 2a**) and accompanying t-SNE and UMAP plots (**Figures 2b-c**), certain biomarkers – indicative of specific cell types - readily grouped with one another and revealed 11 distinct phenotypic clusters. The cellular identity of each cluster was subsequently assigned based on the expression profiles of the 33 markers in each segmented cell. Monocytes, M2 macrophages, CD8 T cells, CD4 T cells, proliferating cells, natural killer cells, dendritic cells, and M1 macrophages were identified as having high expression of CD11b, CD163, CD8, CD4, Ki67, CD57, CD11c, and CD68 respectively. B cells were identified by expression of cell lineage markers, namely CD19, CD20, CD21, and Pax5 which is involved in B cell development and function. Additionally, epithelial (TP63, CD227, CK17, β-catenin, E-cadherin, CK19, and PanCytokeratin) and vascular (Collagen IV, CD31, SMA, CD34) clusters were annotated based on the expression of known markers for each phenotype. The overall abundance and distribution of the 11 phenotypes demonstrates high percentages of B (22%) and T cells (28%), which is consistent with the immune function of tonsil tissue (**Figure 2b-c Legend**). Each phenotyped cell can also be mapped back on to the xy coordinates of the tissue, thereby revealing its spatial coordinates and localization relative to other cells; in this way, we generated a spatial phenotypic map of the cellular organization of the tonsil tissue at single cell resolution (**Figure 2d**).

After targeted spatial phenotyping of tonsil tissue into 11 major phenotypes, we extended our unsupervised clustering to additional markers in the panel. The marker correlation dendrogram in **Figure 2e** shows correlations between immune, cell lineage, metabolic, proliferation, stress, and structural markers in the panel. Markers that co-expressed in certain cell types grouped adjacent to one another, resulting in distinct expression matrices where related molecules indicated the *type and state* of a particular cell. For example, ZAP70, a tyrosine kinase involved in T cell development and lymphocyte activation was found to colocalize with T cell lineage markers CD3, CD7, CD4, and CD2 in the bottom right corner of the dendrogram (**Figure 2e**). This observation could also be confirmed with single-cell resolution anatomical data, as is shown in the co-localization of ZAP70 to T cells in tonsil tissue (**Figure 2f**). Similarly, PAX5, a critical transcription factor for the commitment of lymphoid progenitors to the B cell lineage was highly expressed in CD20+ CD21+ B cells along with CD40 and HLA-DR (**Figure 2e**, **g**). Epithelial, myeloid, macrophage, and vascular markers were also found to associate together in distinct clusters, thus providing further verification of our panel content. Overall, analysis of the multiplex panel of immune and structural markers in human tonsil tissue revealed 11 distinct cellular phenotypes. These phenotypes can be further expanded upon and classified based on expression of metabolic, proliferation and stress markers to get deeper insights into the spatial, cellular, and functional heterogeneity within this tissue.

### Spatial *immune* phenotyping of a human head and neck cancer reveals 6 distinct cellular neighborhoods (CN)

**Figure 3a** shows the H&E-based pathological annotations for a HNSCC clinical tissue sample. The patient, a 55-year-old male, developed recurrent pulmonary and nodal metastatic disease two years after initial surgery and post-operative chemoradiotherapy with concurrent cisplatin for cT3N2M0 p16+ tonsil SCC. Upon relapse, he was treated with 13 cycles of pembrolizumab and showed a partial response in his measurable disease. Treatment was ceased early due to the development of a grade 3 immune-related polyarthritis requiring oral corticosteroids and methotrexate with good effect. His disease then developed subsequent progression approximately 14 months after commencing immunotherapy, requiring palliative chemotherapy (carboplatin and 5-flurouracil). After an initial response to chemotherapy, his disease progressed further, and the patient died approximately 33 months after starting immunotherapy.

The H&E-based histological examination **(Figure 3a)** of the patient’s tissue based on eosinophilic and basophilic affinities revealed valuable morphological information including a normal tonsil region with germinal centers, esophageal submucosal glands (ESMG) and dense tumor cell proliferation; however, it tells us little about the biological processes that underlie the cancer response and progression. Tumorigenesis is complex and highly dynamic and requires the analysis of several biomarkers in order to generate a mechanistic understanding. To address this complexity, we applied our 101-plex antibody panel to this HNSCC tissue to analyze key hallmarks of cancer pathogenesis. StarDist segmentation identified 856,676 cells and we analyzed each of these for expression of 101 protein biomarkers. A representative image (**Figure 3b**) showing epithelial (E-cadherin), stromal (Vimentin), vascular (CD34) and immune cells (CD21 B cells, CD4 T cells) provides an overview of the tumor landscape.

Unsupervised clustering and spatial phenotyping with 33 immune and structural biomarkers, as was initially done in the tonsil (see **Figure 2**), allowed us to annotate 14 cell phenotypes, including immune cell subsets, vasculature, esophageal submucosal glands (ESMG), and lymphatic ducts (**Figure 3c-e**). A cluster expressing epithelial markers along with high expression of PD-L1 and PCNA was also found and annotated as tumor cells (**Figure 3c-e**). Quantification of cell phenotypes revealed high percentages of tumor cells (27%), vasculature (17%), B cells (11%), CD4+ (13%), and CD8+ T cells (8%) relative to other cell types. The spatial map (**Figure 3f**) shows the single-cell geographical distribution of all 14 cell types and reveals the complex architecture of the tissue sample, which included large tumor regions (annotated T in **Figure 3f**) as well as a relatively normal lymphoid region containing tonsillar germinal centers (NT in **Figure 3f**). The marker correlation dendrogram in **Figure 3g** includes the full clustering analysis of biomarkers in our panel and shows groupings of markers that labeled the same cell type(s). Similar patterns emerged as in the tonsil tissue, with distinct matrices related to epithelial, vascular, and immune subsets. A plethora of additional correlations highlight the complexity of the tumor biology and warrant further investigation beyond the scope of this manuscript.

Tissues are organized into complex cellular neighborhoods (7,8) that orchestrate biological processes involved in maintaining tissue homeostasis in health and disease. Crucially, CNs can be identified by *unsupervised* bioinformatic analyses and thus provide an *unbiased* regional breakdown of a tissue’s structure-function relationship. Our CN analysis of the 14 cellular phenotypes described above revealed 6 unique cellular neighborhoods, including CN1 enriched in tumor cells (78%); CN2 enriched in vasculature and lymphatics (81%); CN3 with an abundance of CD4+ T cells (63%); CN4, which consisted almost exclusively of B cells (87%); CN5 with a mixture of CD4+ T cells (11%), CD8+ T cells (17%), macrophages (17%) and tumor cells (12%) as well as CN6 consisting mainly of normal squamous epithelia (91%) (**Figure 4a-b**). The spatial distribution of the CNs across the whole tissue section is shown in **Figure 4c** and corroborates well with the anatomical and cell phenotyping data shown in **Figure 3**. For example, CN3 and CN4 consist mainly of CD4+ T cells and B cells, respectively, and represent the regional morphology of healthy tonsil tissue, consistent with known tonsillar morphology. Interestingly, several small CN4 regions are also located within the tumor tissues and likely representative of tertiary lymphoid structures (see TLS in **Figure 4c-d**). Another neighborhood, CN5 contains a sizeable population of CD8+ T cells and macrophages (see plot in **Figure 4a**), which is consistent with the visible infiltration of M1 macrophages (green) and CD8+ T cells (cyan) seen in the single cell phenotyping map (**Figure 3f**). Overall, this result demonstrates the acuity of unsupervised cellular neighborhood analyses for obtaining objective compositional analyses of complex tissue samples.

**Figure 4:**
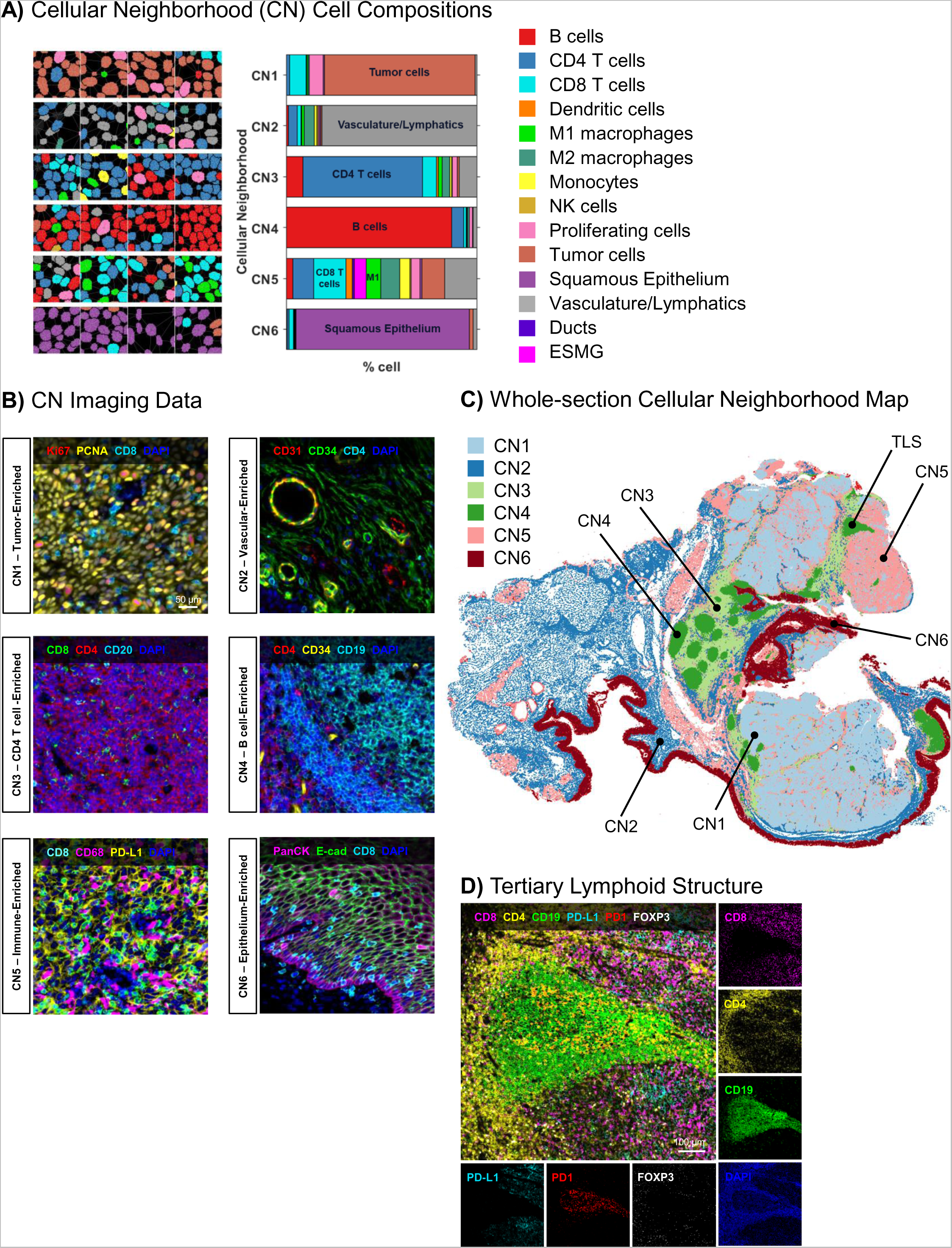
Single-cell spatial phenotyping of a human FFPE head and neck squamous cell carcinoma reveals 6 spatial neighborhoods across 14 distinct cell types. A) Spatial immune phenotyping and cellular neighborhood (CN) analysis of all 0.8 million cells of the tissue into 14 distinct cell types (color code shown in the legend on the right) and 6 cellular neighborhoods: CN1 – Tumor-enriched, CN2 – Vascular-enriched, CN3 – CD4 T cell-enriched, CN4 – B cell-enriched, CN5 – Mixed immune-enriched and CN6 – Squamous epithelium-enriched. Examples of CNs are illustrated with cell nuclei color coded with the representative cell type on the left of the stacked bar graph. B) Examples of CNs with cell phenotypic markers as indicated in the legends. C) A cellular neighborhood spatial map shows the distribution of all the CNs across the tissue. D) A tertiary lymphoid structure (TLS) was also observed as part of CN4 within the tumor region. Expression of CD8 (pink), CD4 (yellow), CD19 (green), PD-L1 (cyan), PD1 (red) and FOXP3 (white) shows the distribution of PD-L1+ tumor cells, CD4+ helper T cells, CD8+ cytotoxic T cells and FOXP3+ T regulatory cells around the CD19+ B cells.

**Figure 5:**
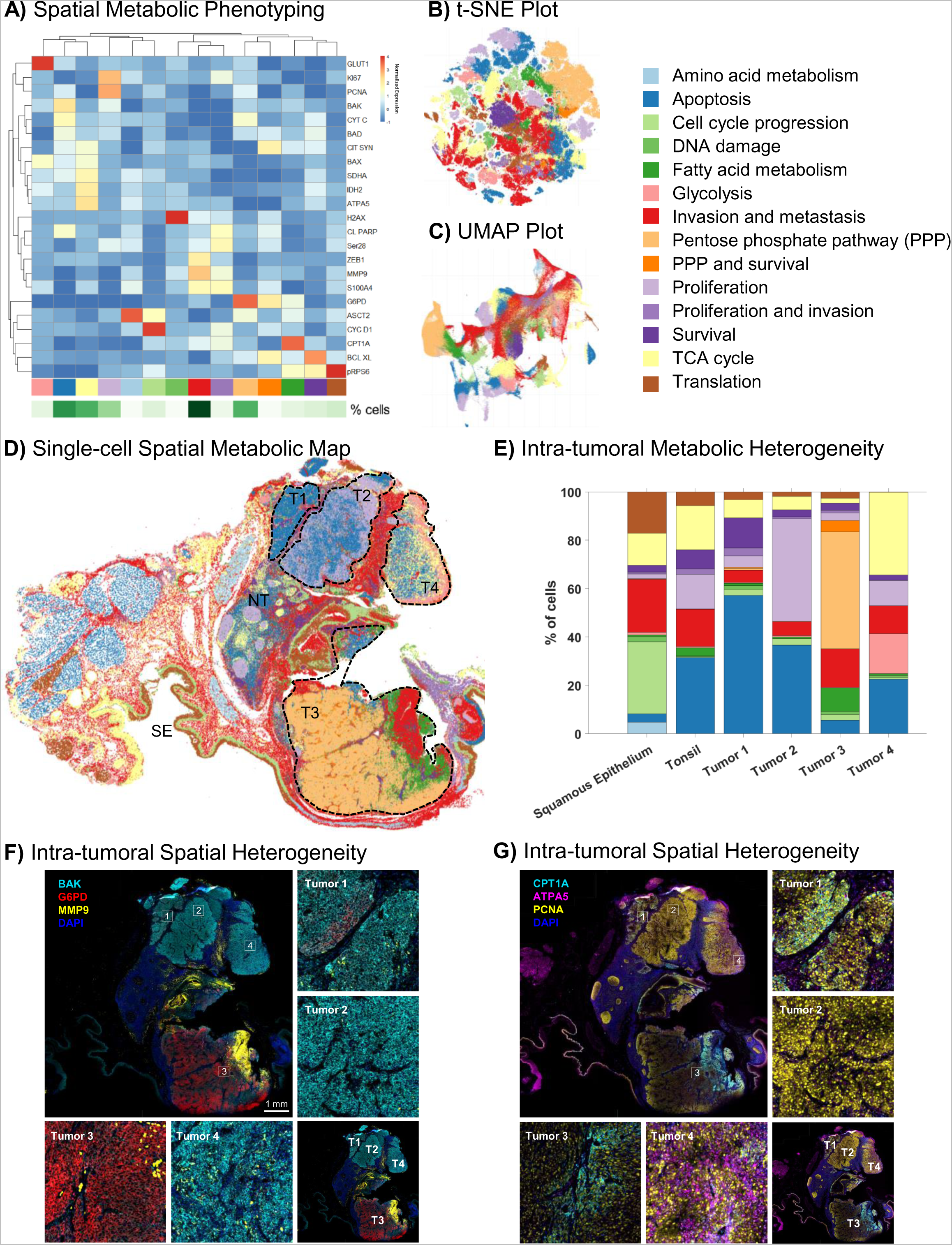
Single-cell spatial metabolic phenotyping of a human FFPE head and neck squamous cell carcinoma reveals 4 distinct tumor regions with high intra-tumoral heterogeneity. A) Unsupervised Leiden clustering and spatial phenotyping based on metabolic and stress markers as indicated on the right of the heatmap identifies 14 distinct functional phenotypes (legend on far right). B) and C) show, respectively, the t-SNE and UMAP plots of all cells colored by metabolic phenotype. D) A spatial metabolic map reveals four distinct tumor regions with varying metabolic and stress profiles. Tumor 1 (T1) has a high degree of apoptotic cells (blue); Tumor 2 (T2) shows a central apoptotic core (blue) surrounded by proliferating cells (light purple); Tumor 3 (T3) shows high expression of markers for the pentose phosphate pathway (peach), fatty acid metabolism (green) and invasion (red); and Tumor 4 (T4) is highly dynamic with cells undergoing glycolysis (light pink), TCA cycle (yellow) and apoptosis (blue). E) Quantification of the 14 functional phenotypes shows the predominant pathways within the normal squamous epithelium, normal tonsil and four tumor regions. F) Intra-tumor spatial heterogeneity in expression of metabolic and stress markers is shown in multiplex images of BAK (cyan), G6PD (red) and MMP9 (yellow) across the four tumor regions. G) Similarly, multiplex images of CPT1A (cyan), ATPA5 (pink) and PCNA (yellow) in the four distinct regions further illustrate the heterogeneity within this sample.

### Moving beyond the immune contexture: *functional metabolic* phenotyping of the head and neck cancer sample reveals additional degrees of intra-tumor heterogeneity

Cellular neighborhoods have gained momentum as a potential prognostic biomarker to stratify immunotherapy responsiveness (7,8). However, evaluation of spatial biomarkers beyond the immune microenvironment has thus far not been attempted. We thus explored the spatial biology of metabolic, proliferation, and stress pathways across the HNSCC tumor to elucidate the functional underpinnings of tumor heterogeneity in greater detail. We used the same analytical logic as above, but instead of focusing on canonical immune biomarkers, we targeted our clustering analyses on 23 markers involved in metabolism, invasion, apoptosis, cell cycle, proliferation, and survival (**Supplementary Figure 4**). Clustering based on these biomarkers allowed us to affirmatively annotate 14 unique *functional* cell phenotypes (**Figure 5a-c)**. The spatial map (**Figure 5d**) resulting from this analysis revealed added complexity and heterogeneity when compared to the neighborhood map obtained based on our analysis of immune markers only (compare **Figures 4c** vs. **5d**). Thus, while spatial phenotyping and cellular neighborhood analysis based on immune and structural markers revealed 6 CNs within the tissue sample (**Figure 4c**), our analysis of metabolic and functional phenotypes revealed a more nuanced functional profiling across all tissue substrates that are contained within the sample (**Figure 5d**). Based on spatial metabolic phenotyping, we therefore estimated that the tumor mass consists of 4 distinct regions. Tumor regions 1, 2, and 4 showed signs of apoptosis based on relatively higher expression of BAK, a pro-apoptotic regulator (**Figure 5f**); all three regions were separated from one another by a distinct margin of immune and invasive cells (see **Figures 3f and 5d**). Tumor region 2 also showed unique spatial features with a central core of apoptotic cells surrounded by a rim of proliferating cells (**Figure 5d**). On the other hand, tumor region 3 showed high levels of invasion as seen by higher expression of Glucose- 6-Phosphate Dehydrogenase (G6PD) and MMP9 (**Figure 5d, f**). G6PD is an enzyme involved in the pentose phosphate pathway and reported to be a mediator of clinical resistance (30,31). Together with carnitine palmitoyltransferase 1A (CPT1A), a rate-limiting enzyme in fatty acid metabolism implicated in metabolic reprogramming for cancer pathogenesis (32), our data in **Figure 5d and f** suggest that tumor region 3 represents a more active tumor phenotype, one that is actively involved in tumor promotion and metastasis relative to the rest of the tumor mass. Additionally, elevated expression of ATPA5, an ATP synthase, in Tumor region 4 (**Figure 5g**) along with pro-apoptotic and immune markers suggests that this region is primed with the energy requirements for apoptosis and immune activation.

In a final analysis, we revisited the distribution of immune-related cell phenotypes (**Figure 3c**) within the four metabolically distinct tumor regions identified above. Since this sample also contained a small region with intact tonsil architecture, it allowed us to compare the cellular distributions in normal tonsil tissue versus the deviation that occurs with tumorigenesis. The normal tonsil region had predominantly B cells (36%), T cells (41%), and vasculature (16%) (**Figure 6a**). In contrast, the tumor regions had a high abundance of proliferating tumor cells (25% to 63%) with relatively few B cells. Interestingly, tumor region 4 had a large B cell cluster, which displayed multiple morphological features that are consistent with tertiary lymphoid structures (see **Figure 4d**). Tumor region 4 also showed high levels of CD68+ M1 macrophages (18%) and CD8+ T (28%) cells, suggesting high anti-tumor immune activity in this region (**Figure 6a**). Tumor region 3, on the other hand, had higher levels of CD163+ M2 tumor-associated macrophages (3%) relative to M1 macrophages (0.4%; **Figure 6b**). Spatial proximity analyses of the four distinct tumor regions further revealed that M1 macrophages were found to be closely associated with tumor cells in tumor region 4 while M2 macrophages were found to be farther away (**Figure 6c**). This trend is reversed with a closer association of M2 macrophages relative to M1 macrophages with tumor cells in tumor region 3. These results further highlight the importance of single-cell spatial phenotyping and analyzing cellular interactions within the context of the tissue architecture to understand the complex nuances of the tumor microenvironment.

**Figure 6:**
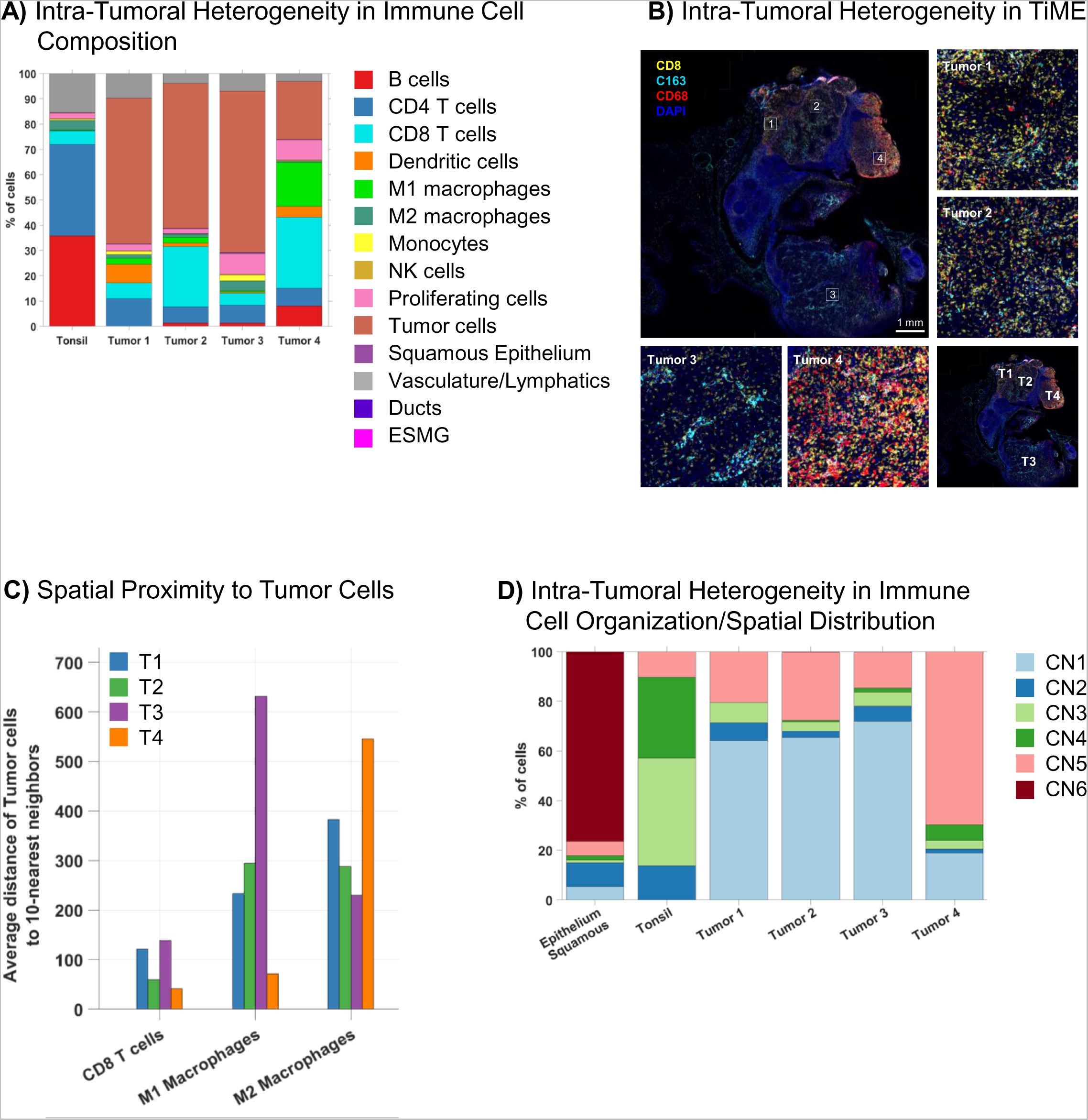
Differences in immune cell composition and spatial neighborhood distribution within the 4 distinct tumor regions reveal diverse tumor immune microenvironments within the same sample. A) Quantification of the relative abundance of 14 cellular phenotypes (legend on the right) in the normal tonsil region and the four distinct tumor regions identified based on metabolic phenotyping shows higher abundance of CD4+ T cells (blue), B cells (red) and vasculature (gray) in tonsil relative to tumors. Notable differences are also observed in CD8+ T cells (cyan) and M1 macrophages (light green) compositions between the four tumor regions with a higher percentage of CD8+ T cells in Tumor region 2 and 4 and a higher percentage of M1 macrophages in Tumor region 4. B) Multiplex images of CD8 (yellow, cytotoxic T cell marker), CD163 (cyan, M2 macrophage marker), CD68 (red, M1 macrophage marker) and DAPI (blue) shows the spatial heterogeneity of CD8+ T cells and macrophages within the four tumor regions and highlights the different tumor immune microenvironments in a single HNSCC sample. C) Spatial proximity analysis further quantifies differences in the distances between tumor cells and immune cells (CD8 T cells, M1 macrophages and M2 macrophages) within the four tumor regions. CD8+ T cells and M1 macrophages are closer to tumor cells in Tumor region 4 (orange) relative to Tumor region 3 (purple); M2 macrophages are closer to tumor cells in Tumor region 3 (purple) relative to Tumor region 4 (orange). D) Differences in the abundance of cellular neighborhoods identified in Figure 4 suggest a higher order organization of immune and other (tumor, epithelial and vascular) cells in normal tonsil and tumor regions. Tumor regions 1, 2 and 3 have similar macro-organization of cellular neighborhoods with a higher percentage of CN1 (tumor-enriched) relative to Tumor 4 which has a higher percentage of CN5 (immune-enriched). Abundance of CN2 (Vascular-enriched), CN3 (CD4 T cell-enriched), CN4 (B-cell enriched) and CN6 (epithelium-enriched) is also shown here.

Lastly, we re-analyzed the abundance of cellular neighborhoods within the four tumor regions vs. adjacent healthy tonsil. The distribution of the CNs within the four tumor regions revealed differences in spatial organization of immune-centric cells at a macro level (**Figure 6d**). A higher percentage of CN5 (mixed immune neighborhood) (69%) was observed in Tumor region 4 relative to the other three tumor regions (T1 - 20%, T2 - 28% and T3 - 15%). Tumor region 4 also had a lower overall tumor burden with lower levels of CN1 (tumor-rich neighborhood; 18%) compared to Tumors 1 (63%), 2 (65%), and 3 (72%). The degree of intra-tumoral heterogeneity within this tissue section was also apparent by differences in the levels of CN3 (CD4-rich neighborhood) between the 4 tumor regions, with Tumor region 1 having a slightly higher percentage of CN3 (8%) compared to the other three regions (T2 - 3%, T3 - 6% and T4 - 2.5%). Overall, these data amount to a uniquely comprehensive account of the structural and functional topography of a HNSCC tissue sample. In doing so, our data highlight the potential utility of in-depth analysis of spatial phenotyping as a tool for clinical discoveries.

## DISCUSSION

Head and neck squamous cell carcinomas (HNSCC) remain an ongoing global challenge to public health (1). Despite the approval of immunotherapies to treat metastatic and recurrent HNSCC, the overall survival rate remains low, partially due to heterogenous tumor microenvironments and plasticity of the HNSCC tumor immune landscape (3). With molecular classification of HNSCC tumors and integrated genomic and epigenetics data from TCGA, new insights into the molecular and genetic landscape of HNSCC have now emerged (33,34). However, these studies lack spatial context and/or an evaluation of the tumor immune microenvironment at the functional level. Understanding the spatial context of the tumor microenvironment as well as key hallmarks of HNSCC pathogenesis, progression and resistance will be a critical next step towards elucidating the complex tumor biology and mechanistic principles that underlie differential clinical outcomes.

### Decoding HNSCC Tumor Biology: Towards a holistic view of HNSCC tumor biology

Unsupervised spatial phenotyping of a patient’s HNSCC sample allowed us to reconstruct the patient’s tumor immune microenvironment with unprecedented detail. Starting with the identification of a dozen immune-related cell phenotypes throughout the tissue (**Figure 3f**) to unsupervised classification of immune cell neighborhoods (**Figure 4c**), our analysis provided a detailed spatial topography of the tissue. Differences in immune infiltration across the tissue were immediately evident along with the presence of tertiary lymphoid structures (TLS) (**Figure 4c-d**). All of these findings relate to immunotherapy outcomes and hence harbor analytical value. For example, tertiary lymphoid structures are a type of cellular neighborhood comprised of a distinct organization of B cells, T cells, dendritic cells and high endothelial venules (35,36). TLS have been reported to play a role in the anti-tumor response across multiple cancer types (37–40). The presence of TLS in most cases is associated with better prognosis, and, hence, they have emerged as a valuable predictor of response to ICI therapy (39–42). The presence of the TLS, in the HNSCC tumor described here, in a region marked by infiltration of CD8+ and CD4+ T cells points to an immunologically active microenvironment in the top right of the tissue (**Figure 4c-d**). Understanding the spatial organization and composition of TLS in the microenvironment of HNSCC, including the formation and dynamics of TLS through disease progression and treatment will be important to understand more about the role of these structures in anti-tumor responses.

Many additional biological processes affect tumor progression and treatment responses beyond the tumor immune microenvironment. For example, deregulation of cellular energetics and metabolic reprogramming is a key hallmark of cancer. While most studies focus on mass spectrometry or extracellular flux analysis for quantification of cellular metabolites, these studies lack single cell resolution and spatial information. A single-cell spatial metabolic profiling study of cytotoxic T cells in colorectal cancer specimens identified cells in unique tumor-specific metabolic states associated with distinct immune phenotypes and demonstrated the power of combining metabolic analysis with spatial resolution (43). Leveraging expression data from 23 proteins involved in critical metabolic and stress pathways in tumors, we developed a spatial map of key cancer metabolism targets illustrating the predominant functional pathways in the HNSCC tumor. Functional mapping across every cell within the entire tissue section led to deeper stratification and identification of distinct regions in the tumor with unique metabolic and stress profiles. Tumor regions 2 and 4, with high immune cell infiltration based on spatial immune phenotyping (**Figure 3f**) also showed high expression of apoptotic markers (**Figure 5d-f**), suggesting an active anti-tumor immune response in this region. On the other hand, within the same tumor, Tumor region 3 with densely packed tumor cells had high expression of markers for DNA damage, invasion, and pentose phosphate pathway. Overexpression of G6PD has been reported in several cancers (44,45) and is believed to mediate protection against oxidative stress. Moreover, increased levels of G6PD are associated with resistance and poor prognosis in the clinic. Our data reveal the high degree of intra-tumoral heterogeneity with pro-tumorigenic and anti-tumor immunologically active regions in the same tissue. Combining phenotypic lineage information with metabolic and stress markers provides information (**Supplementary Figure 5a-b**) on the key functional determinants of specific immune subsets and tumor cells. This can aid in understanding the key mechanisms in dynamic and evolving tumor microenvironments that contribute to the overall clinical response to immune therapies and in identifying new targets for their regulation.

### Spatial Biomarkers and their Translational Relevance

H&E staining is routinely used in clinical practice to diagnose and classify (46,47) cancers. The H&E-based pathological assessment of the HNSCC tissue sample studied here revealed a normal tonsil region and a large tumor (**Figure 3a**). While histological examination of tissues based on eosinophilic and basophilic affinities of tissue structures can reveal valuable morphological information, they are limited in biomarker depth and do little to inform us about cellular phenotypes or functions. In contrast, the data that we have obtained via antibody-based spatial phenotyping to detect protein expression and localization show a far greater depth of information than what is available via H&E staining, and they enable the exciting possibility of generating a holistic view of the tumor microenvironment and key cancer hallmarks *in situ* at single-cell resolution.

Indeed, several recent studies have highlighted the value of highly multiplexed spatial phenotyping of the tumor immune microenvironment for stratification of responses to immunotherapy. A study in colorectal cancer (8), for example, introduced a novel analysis framework – cellular neighborhood analysis - to analyze the organizational patterns of distinct immune cell types in the tumor. Applying a 56-plex antibody panel across a cohort of advanced colorectal cancer tissues, the study revealed the correlation between clinical outcomes and distinct organizational patterns including neighborhoods enriched for tertiary lymphoid structures, T cells, macrophages, and granulocytes. In a similar retrospective study on cutaneous T cell lymphoma (7), researchers sought to identify differences in responsive and non-responsive cohorts following treatment with pembrolizumab. Single-plex IHC studies for CD4, PD1, PD-L1 and FOXP3 revealed no significant differences between the two groups; neither did an analysis of the tumor mutational burden. However, with a 55-plex CODEX panel, the authors found a higher frequency of CD4+ immune-activated CNs in responders vs. non-responders. These findings were accompanied by significantly higher levels of Treg-enriched immune-suppressed CNs in the non-responding cohort before and after pembrolizumab treatment. Thus, in contrast to several well-established clinical diagnostic biomarkers that failed to reveal a difference between responding and non-responding patients, CNs represent a new class of spatial biomarkers that harbor the possibility to better stratify immunotherapy patient cohorts. These types of findings emphasize the translational significance of high-plex biomarker discoveries to derive predictive signatures of clinical relevance.

### Challenges and Future Directions

An ongoing challenge to the widespread deployment of high-plex spatial phenotyping technologies is the tradeoff between imaging resolution limits, biomarker plex level and data collection speed. To our knowledge, ours is the first study to demonstrate *in-situ* analysis of >100 proteins via optical imaging. Juxtaposing the H&E map (**Figure 3a**) with the single-cell spatial phenotypic map (**Figure 3f**), the whole-section immune cellular neighborhood map (**Figure 4c**) and the metabolic functional map (**Figure 5d**) unveils the different layers of complexity of the tumor microenvironment. The sheer complexity and depth of these single-cell spatial and functional data raises several hypotheses for the mechanisms underlying partial clinical responses and provides guiding principles for the application of such high-plexed discovery-based research tools to identify biomarkers of clinical significance.

Further studies are warranted where this approach is used to screen larger cohorts of patients with pathological complete responses and progressive disease to distill multi-marker and multi-omic spatial signatures associated with therapy response and resistance. These tissue signatures can then be prospectively validated and developed into screening tools/companion diagnostic assays for predicting response to immunotherapies in head and neck cancer.

## Acknowledgements

The authors thank The Passe and Williams Foundation for the grant support issued to AK and BGMH.

