## Supplemental Information for "Single-cell Spatial Metabolic and Immune Phenotyping of Head and Neck Cancer Tissues Identifies Tissue Signatures of Response and Resistance to Immunotherapy"

### SUPPLEMENTARY INFORMATION

#### SUPPLEMENTARY FIGURE LEGENDS

**Supplementary Figure 1:** PhenoCycler staining, imaging and analysis workflows. A) The PhenoCycler workflow enables staining with a multiplex panel of oligo-conjugated antibodies with unique DNA barcodes, followed by iterative steps of hybridization with complementary oligo-tagged fluorescent dyes; three markers + DAPI are subsequently imaged at a time, followed by dehybridization and additional cycles. The reveal-image-remove cycles are automated on the PhenoCycler-Fusion. B) Computational analysis workflow for spatial phenotyping includes a qualitative quality control (QC) assessment of each marker, followed by cell segmentation based on nuclear DAPI staining. Average marker intensities are calculated on a per cell basis using nuclear and membrane masks and unsupervised Leiden clustering is performed for immune and metabolic phenotyping. Clusters are manually annotated based on biomarker expression profiles. Additionally, cellular neighborhood or compartmental analyses may be performed to provide the spatial organization of different cell types on a cellular and regional scale, respectively.

**Supplementary Figure 2:** Cellular neighborhood analysis. A) Delaunay Triangulation is used to compute cellular neighborhoods. A cell 'N' is considered a neighbor of a cell 'A' if it is connected in terms of a Delaunay triangulation. B) Based on Calinski-Harabasz criteria, the optimal number of cellular neighborhoods was determined to be 6. C) Average neighboring cell-type percentages are calculated for all cells in each cellular neighborhood.

**Supplementary Figure 3:** Validation of CODEX antibodies. A) Comparison of CODEX antibody performance on PhenoCycler Fusion (left panel) with orthogonal DAB IHC (right panel) staining for PD-1, PD-L1 and FOXP3 shows comparable staining and expression patterns in human tonsil tissue. B) Representative images of all 101 markers in the panel demonstrating high sensitivity and specificity in human FFPE tissues.

**Supplementary Figure 4:** The 101-plex panel comprises of 5 modules for (1) tumor-immune phenotyping and neighborhood mapping, (2) proliferation, stress and death profiling, (3) mapping of tumor structure, invasion and metastasis, (4) signaling pathway analysis and (5) metabolic phenotyping of human FFPE tissues. Each module reveals valuable information on one or more hallmarks of cancer such as immune evasion, cellular energetics, angiogenesis, invasion and metastasis, sustained proliferation and resisting cell death. When combined together in a multiplex panel, it can provide critical information on the intersecting predictors of tumor pathogenesis and progression.

**Supplementary Figure 5:** A) Unsupervised Leiden clustering and heatmaps showing the differential expression of metabolic, immune and stress markers in specific immune (CD4+ T cells, CD8+ T cells, M1 macrophages, M2 macrophages) and epithelial/tumor cell subsets within normal tonsil (NT), normal squamous epithelia (SE) and the four distinct tumor regions as identified by functional metabolic phenotyping. B) Expression profiles of key functional markers in the 14 cellular phenotypes (legend on the right) across the HNSCC tissue shows the predominant metabolic and stress pathways in each cell type.

**Supplementary Table 1: List of Antibodies in the 101-Plex Panel**

| ***Target*** | ***Clone*** | ***Dilution*** | ***Vendor*** |
|---|---|---|---|
| ASCT2 | CAL33 | 1:50 | Abcam |
| ATPA5 | EPR13030 (B) | 1:50 | Abcam |
| AXL | EPR19880 | 1:50 | Abcam |
| β-actin | AKYP0072 | 1:50 | Akoya (Cat. No. 240201) |
| BAD | Y208 | 1:50 | Abcam |
| BAK | Y164 | 1:50 | Abcam |
| BAX | E63 | 1:200 | Abcam |
| β-catenin | AKYP0068 | 1:100 | Akoya (Cat. No. 240200) |
| BCL-XL | E18 | 1:50 | Abcam |
| BECLIN1 | EPR20473 | 1:50 | Abcam |
| C1QA | EPR2980Y | 1:50 | Abcam |
| Caveolin | AKYP0115 | 1:200 | Akoya (Cat. No. 240183) |
| CCR6 | EPR22259 | 1:50 | Abcam |
| CD107a | AKYP0004 | 1:200 | Akoya (Cat. No. 232125) |
| CD11b | AKYP0087 | 1:200 | Akoya (Custom) |
| CD11c | AKYP0051 | 1:200 | Akoya (Cat. No. 232177) |
| CD138 | EPR6454 | 1:50 | Abcam |
| CD14 | AKYP0079 | 1:50 | Akoya (Cat. No. 240066) |
| CD141 | AKYP0124 | 1:200 | Akoya (Cat. No. 240192) |
| CD15 | HI98 | 1:500 | BioLegend |
| CD163 | D6U1JT | 1:50 | CST |
| CD19 | RM332 | 1:50 | RevMab |
| CD1a | EPR3622 | 1:50 | Abcam |
| CD2 | EPR6451 | 1:50 | Abcam |
| CD20 | AKYP0049 | 1:200 | Akoya (Cat. No. 232175) |
| CD21 | AKYP0061 | 1:200 | Akoya (Cat. No. 240003) |
| CD227/MUC1 | 16A | 1:50 | BioLegend |
| CD31 | AKYP0047 | 1:200 | Akoya (Cat. No. 232172) |
| CD34 | AKYP0088 | 1:50 | Akoya (Cat. No. 240076) |
| CD38 | AKYP0110 | 1:200 | Akoya (Cat. No. 240178) |
| CD39 | AKYP0107 | 1:50 | Akoya (Cat. No. 240175) |
| CD3e | AKYP0062 | 1:200 | Akoya (Cat. No. 240006) |
| CD4 | AKYP0048 | 1:200 | Akoya (Cat. No. 232174) |
| CD40 | AKYP0095 | 1:200 | Akoya (Cat. No. 240083) |
| CD44 | AKYP0073 | 1:200 | Akoya (Cat. No. 232124) |
| CD45 | AKYP0074 | 1:200 | Akoya (Cat. No. 240060) |
| CD45RO | AKYP0059 | 1:50 | Akoya (Cat. No. 232188) |
| CD56 | AKYP0118 | 1:50 | Akoya (Cat. No. 240186) |
| CD57 | HNK-1 | 1:50 | BioLegend |
| CD68 | AKYP0050 | 1:200 | Akoya (Cat. No. 232176) |
| CD69 | EPR21814 | 1:50 | Abcam |
| CD7 | EPR4242 | 1:50 | Abcam |
| CD79a | AKYP0109 | 1:200 | Akoya (Cat. No. 240177) |
| CD8 | AKYP0028 | 1:200 | Akoya (Cat. No. 232151) |

| | | | |
|---|---|---|---|
| Citrate Synthase | EPR8067 | 1:50 | Abcam |
| CK17 | W16131A | 1:100 | BioLegend |
| CK19 | A53/B-A2 | 1:50 | BioLegend |
| Cleaved PARP | E51 | 1:50 | Abcam |
| Collagen IV | AKYP0083 | 1:100 | Akoya (Cat. No. 240070) |
| CPT1A | EPR21843-71-2F | 1:50 | Abcam |
| Cyclin D1 | RM241 | 1:50 | RevMab |
| Cytochrome c | EP1326-80-5 | 1:50 | Abcam |
| E-cadherin | AKYP0057 | 1:200 | Akoya (Cat. No. 232185) |
| EpCAM | D9S3P | 1:50 | Akoya (Cat. No. 240187) |
| FOXP3 | AKYP0102 | 1:200 | Akoya (Cat. No. 240170) |
| G6PD | EPR20668 | 1:50 | Abcam |
| Gal9 | 9M1-3 | 1:50 | BioLegend |
| GATA3 | D13C9 | 1:50 | Akoya (Cat. No. 240184) |
| GLUT1 | EPR3915 | 1:200 | Abcam |
| GP100 | HMB45 | 1:50 | Akoya (Cat. No. 240191) |
| Granzyme B | AKYP0086 | 1:100 | Akoya (Cat. No. 240074) |
| HK1 | EPR10134 (B) | 1:50 | Abcam |
| HLA-A | AKYP0078 | 1:50 | Akoya (Cat. No. 240065) |
| HLA-DPB1 | EPR11226 | 1:50 | Abcam |
| HLA-DR | AKYP0063 | 1:50 | Akoya (Cat. No. 240017) |
| HLA-E | AKYP0096 | 1:50 | Akoya (Cat. No. 240084) |
| ICOS | AKYP0090 | 1:50 | Akoya (Cat. No. 240078) |
| IDH2 | EPR7577 | 1:300 | Abcam |
| IDO1 | AKYP0084 | 1:200 | Akoya (Cat. No. 240071) |
| IFNG | AKYP0093 | 1:200 | Akoya (Cat. No. 240081) |
| iNOS | AKYP0104 | 1:50 | Akoya (Cat. No. 240172) |
| Ki67 | AKYP0052 | 1:200 | Akoya (Cat. No. 232179) |
| LAG3 | AKYP0089 | 1:50 | Akoya (Cat. No. 240077) |
| LaminB1 | EPR8985(B) | 1:100 | Abcam |
| LC3B | EPR18709 | 1:50 | Abcam |
| LDHA | EP1566Y | 1:50 | Abcam |
| LEF1 | EPR2029Y | 1:50 | Abcam |
| MC Tryptase | EPR9522 | 1:100 | Abcam |
| MMP9 | EP1254 | 1:50 | Abcam |
| MPO | AKYP0113 | 1:100 | Akoya (Cat. No. 240181) |
| NaKATPase | EP1845Y | 1:50 | Abcam |
| OX40 | Polyclonal | 1:50 | R&D Systems |
| PanCK | AKYP0053 | 1:200 | Akoya (Cat. No. 232180) |
| Pax5 | RM331 | 1:100 | RevMab |
| PCNA | AKYP0085 | 1:200 | Akoya (Cat. No. 240073) |
| PD1 | AKYP0070 | 1:200 | Akoya (Cat. No. 240035) |
| PD-L1 | RM320 | 1:200 | RevMab |
| pH2AX | 2F3 | 1:100 | BioLegend |
| pHH3 | HTA28 | 1:50 | BioLegend |
| pNRF2 | EP1809Y | 1:50 | Abcam |

| | | | |
|---|---|---|---|
| Podoplanin | AKYP0007 | 1:200 | Akoya (Cat. No. 240203) |
| pRPS6 | SP45 | 1:50 | Abcam |
| S100A4 | S100A4 | 1:50 | BioLegend |
| SDHA | EPR9043 (B) | 1:50 | Abcam |
| SMA | AKYP0081 | 1:50 | Akoya (Cat. No. 240068) |
| SOX2 | SP76 | 1:100 | Abcam (Cat. No. 240174) |
| TFAM | 18G102B2E11 | 1:200 | Akoya (Cat. No. 240020) |
| TP63 | W15093A1 | 1:100 | BioLegend |
| Vimentin | AKYP0082 | 1:200 | Akoya (Cat. No. 240069) |
| ZAP70 | RM408 | 1:50 | RevMab |
| ZEB1 | EPR17375 | 1:50 | Abcam |

**Supplementary Table 2: 101-Plex PhenoCycler-Fusion Experiment Design**

| *Cycle* | *Target* | *Reporter* | *Catalog Number* |
|---|---|---|---|
| 1 | Blank | | |
| 2 | CD39 | ATTO550 | 6250025 |
| | FOXP3 | AF647 | 6550004 |
| | ASCT2 | AF750 | Custom |
| 3 | Podoplanin | ATTO550 | 6250035 |
| | CD68 | AF647 | 6350003 |
| | HLA-A | AF750 | 6450002 |
| 4 | iNOS | ATTO550 | 6250006 |
| | IDH2 | AF647 | Custom |
| | PanCK | AF750 | 6450007 |
| 5 | Ki67 | ATTO550 | 6250012 |
| | CD11c | AF647 | 6550018 |
| | CD79a | AF750 | 6450022 |
| 6 | SOX2 | ATTO550 | 6250026 |
| | Collagen IV | Cy5 | 6350010 |
| | Vimentin | AF750 | 6450008 |
| 7 | IFNG | ATTO550 | 6250005 |
| | LAG3 | AF647 | 6550002 |
| | CD326/EpCAM | AF750 | 6450025 |
| 8 | Granzyme B | ATTO550 | 6250011 |
| | CD40 | AF647 | 6550007 |
| | MC Tryptase | AF750 | Custom |
| 9 | CD8 | ATTO550 | 6250007 |
| | Histone H3 Phospho (Ser28) | AF647 | Custom |
| | LC3B | AF750 | Custom |

| | | | |
|---|---|---|---|
| | Cleaved PARP | ATTO550 | Custom |
| 10 | BCL-XL | AF647 | Custom |
| | NaKATPase | AF750 | Custom |
| | Cytochrome c | ATTO550 | Custom |
| 11 | BAK | AF647 | Custom |
| | CD227 | AF750 | Custom |
| | LEF1 | ATTO550 | Custom |
| 12 | CD138 | AF647 | Custom |
| | GLUT1 | AF750 | Custom |
| | Cyclin D1 | ATTO550 | 6250031 |
| 13 | OX40 | AF647 | Custom |
| | S100A4 | AF750 | Custom |
| | BAD | ATTO550 | Custom |
| 14 | BAX | AF647 | Custom |
| | CD19 | AF750 | Custom |
| | pNRF2 | ATTO550 | Custom |
| 15 | CPT1A | AF647 | Custom |
| | CD45 | AF750 | Custom |
| | MPO | ATTO550 | 6250029 |
| 16 | CD2 | AF647 | Custom |
| | CD57 | AF750 | 6450023 |
| | CD1a | ATTO550 | Custom |
| 17 | ATPA5 | AF647 | Custom |
| | CD44 | AF750 | Custom |
| | GATA3 | ATTO550 | 6250030 |
| 18 | Citrate Synthase | AF647 | Custom |
| | IDO1 | AF750 | Custom |
| | CD56 | ATTO550 | 6250016 |
| 19 | CD3e | AF647 | 6550024 |
| | SMA | AF750 | 6450005 |
| 20 | HLA-E | ATTO550 | 6250022 |

| | | | |
|---|---|---|---|
| | PD-1 | AF647 | 6550006 |
| | Beta-actin | AF750 | 6450027 |
| 21 | TP63 | AF647 | 6550009 |
| | Caveolin | AF750 | 6450024 |
| | CK17 | ATTO550 | 6250021 |
| 22 | PD-L1 | AF647 | 6550005 |
| | | | 6450001 |
| | CD31 | AF750 | |
| | GP100 | ATTO550 | 6250033 |
| 23 | CD7 | AF647 | Custom |
| | G6PD | AF750 | Custom |
| 24 | CD69 | AF647 | Custom |
| | HLA-DPB1 | AF750 | Custom |
| | PCNA | ATTO550 | Custom |
| 25 | Pax5 | AF647 | 6550010 |
| | CCR6 | AF750 | Custom |
| | CD38 | ATTO550 | 6250028 |
| 26 | AXL | AF647 | Custom |
| | CD15 | AF750 | Custom |
| | CK19 | ATTO550 | 6250027 |
| 27 | CD163 | AF647 | 6550001 |
| | C1Qa | AF750 | Custom |
| 28 | SDHA | AF647 | Custom |
| | CD14 | AF750 | 232047 |
| | LaminB1 | ATTO550 | 6250024 |
| 29 | CD4 | AF647 | 6350001 |
| | CD34 | AF750 | 6450009 |
| 30 | ZEB1 | ATTO550 | Custom |

| | | | |
|---|---|---|---|
| | HK1 | AF647 | Custom |
| | CD45RO | AF750 | Custom |
| | CD141 | ATTO550 | 6250034 |
| 31 | LDHA | AF647 | Custom |
| | HLA-DR | AF750 | Custom |
| 32 | MMP-9 | AF647 | Custom |
| | ICOS | AF750 | Custom |
| 33 | pRPS6 | AF647 | Custom |
| | b-Catenin | AF750 | Custom |
| | Galectin 9 | ATTO550 | 6250010 |
| 34 | CD11b | AF647 | 6550020 |
| | CD20 | AF750 | 6450003 |
| | TFAM | ATTO550 | 6250008 |
| 35 | Beclin-1 | AF647 | Custom |
| | E-cadherin | AF750 | Custom |
| 36 | CD107a | AF647 | 6350002 |
| | ZAP70 | AF750 | Custom |
| 37 | H2A.X | AF647 | 6550012 |
| | CD21 | AF750 | Custom |
| 38 | | Blank | |

### Supplementary Figures

**A)** PhenoCycler Workflow

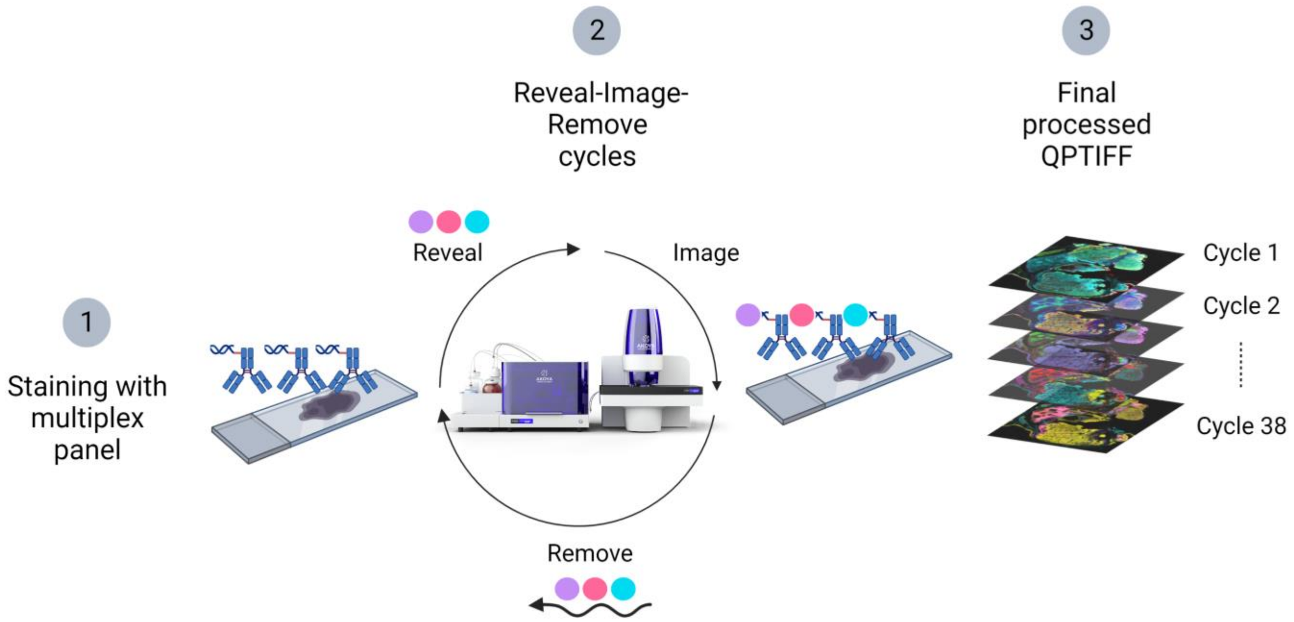


**B)** Bioinformatic Analysis Workflow

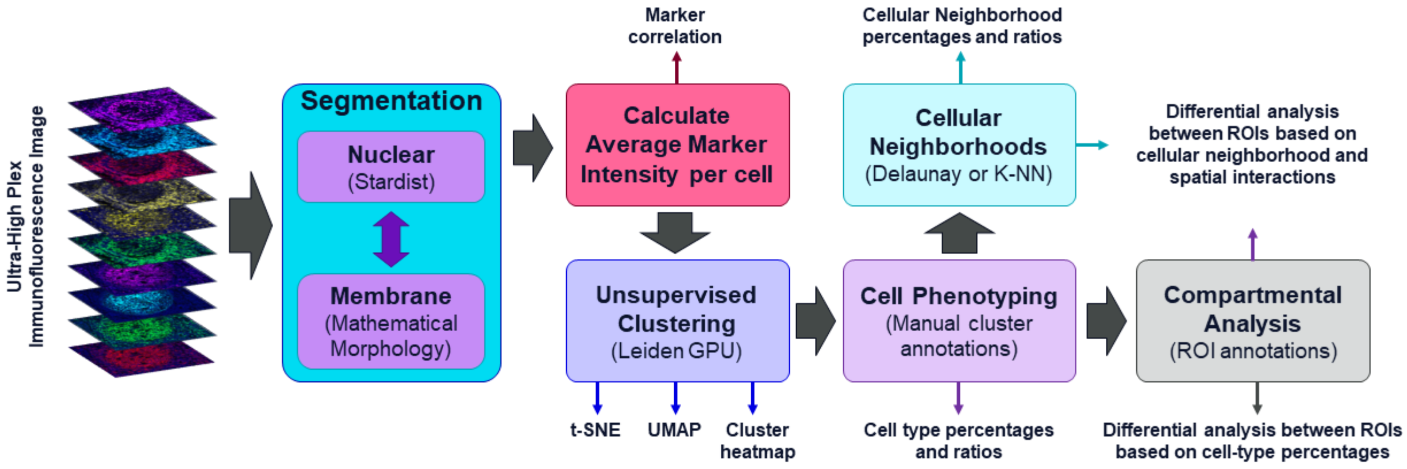


**- Supplementary Figure 1 -**

**A)** Example of Delaunay Triangulation

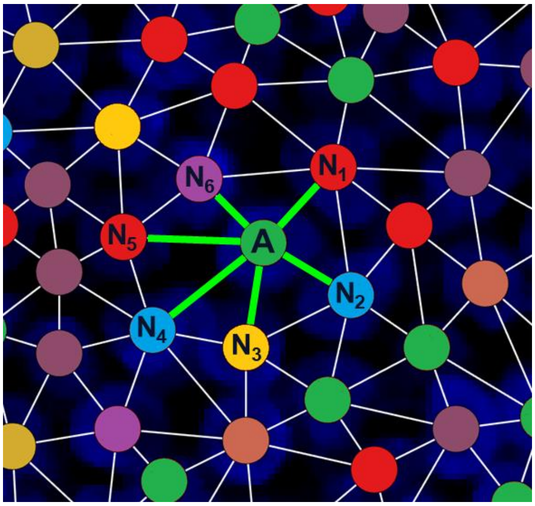


**B)** Calinski-Harabasz criterion

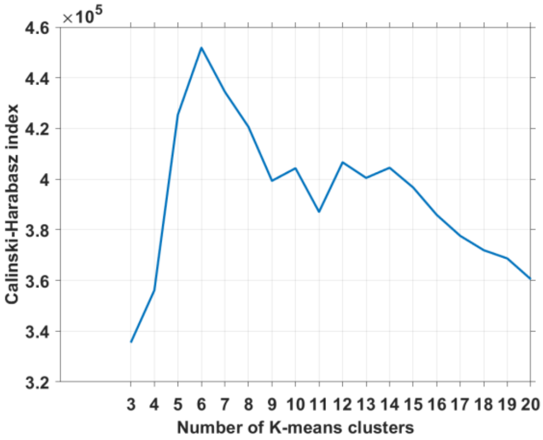


**C)** Percentages of CN per cell type

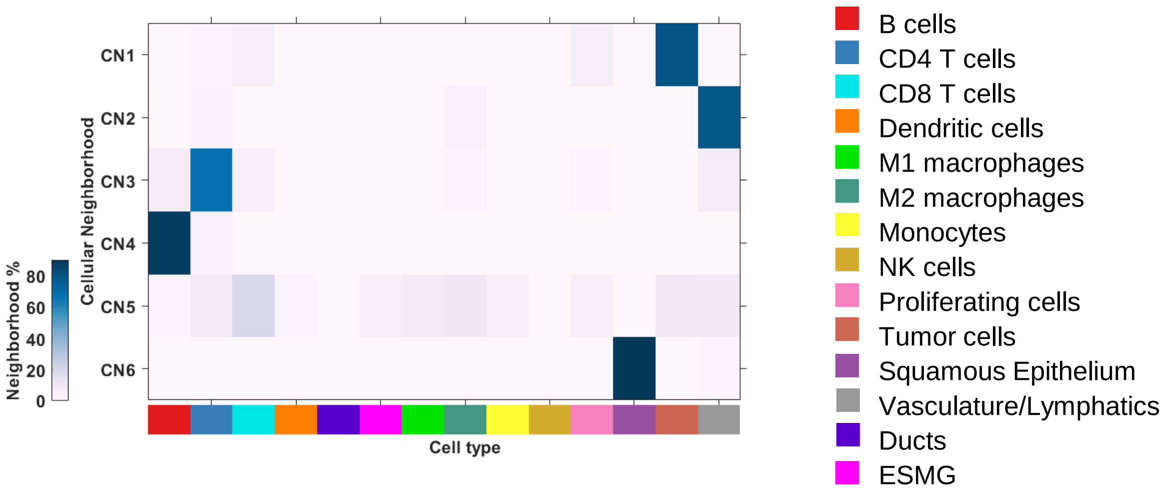


**- Supplementary Figure 2 -**

**A)** DAB IHC vs CODEX PCF Comparison

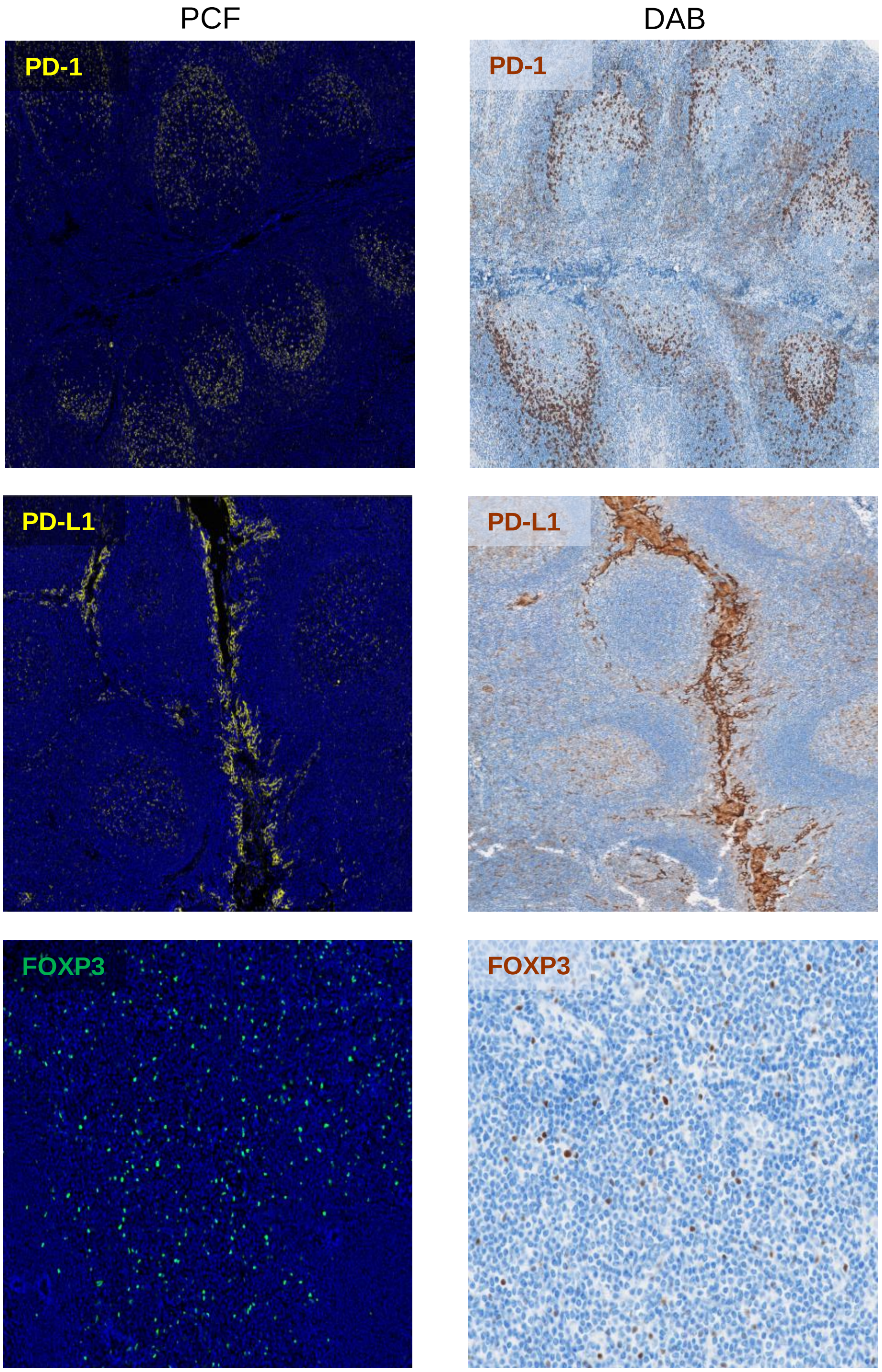


**- Supplementary Figure 3 -**

**B)** Validation of Antibody Expression Pattern

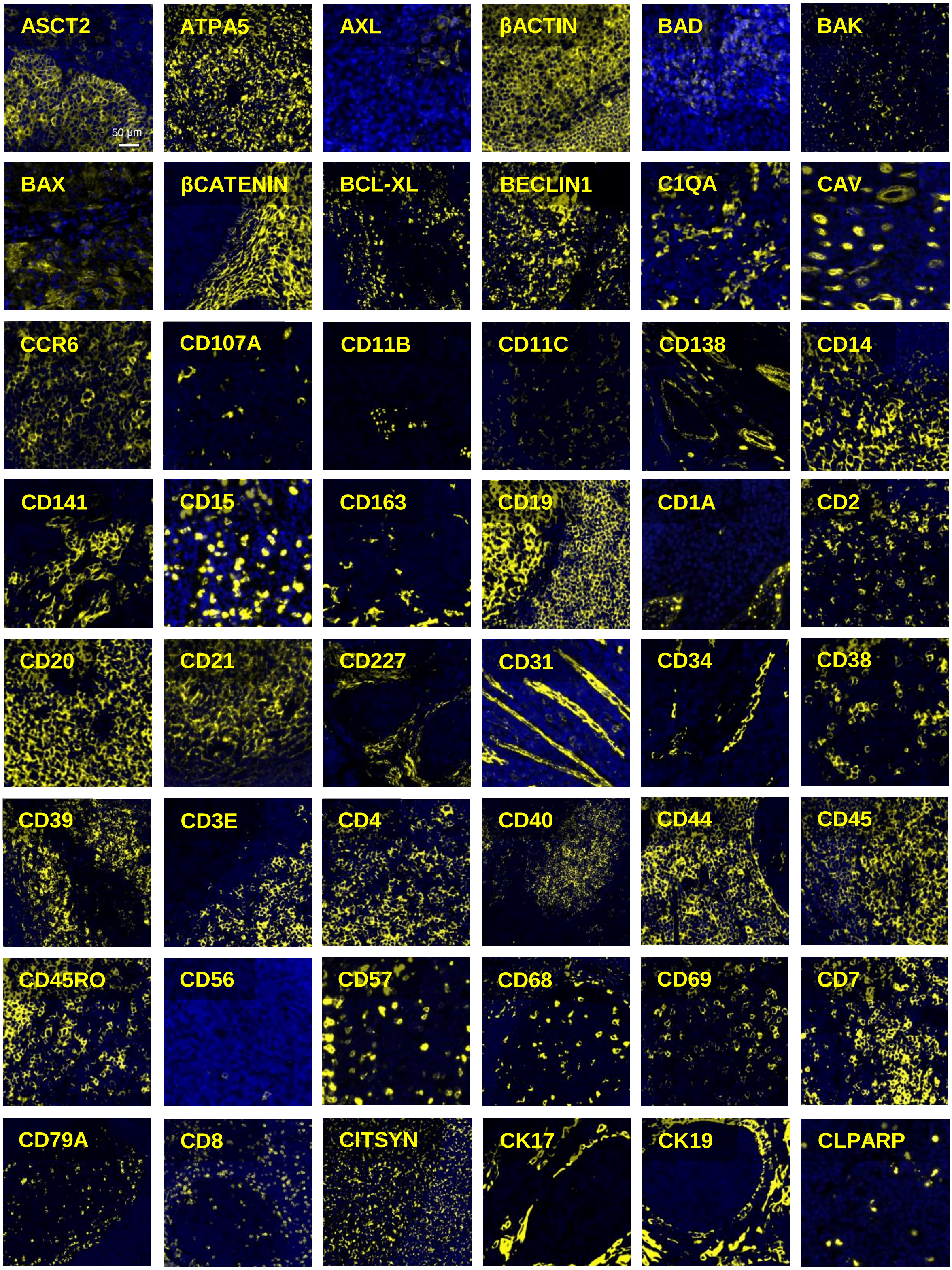


**- Supplementary Figure 3 -**

**B)** Validation of Antibody Expression Pattern (continued)

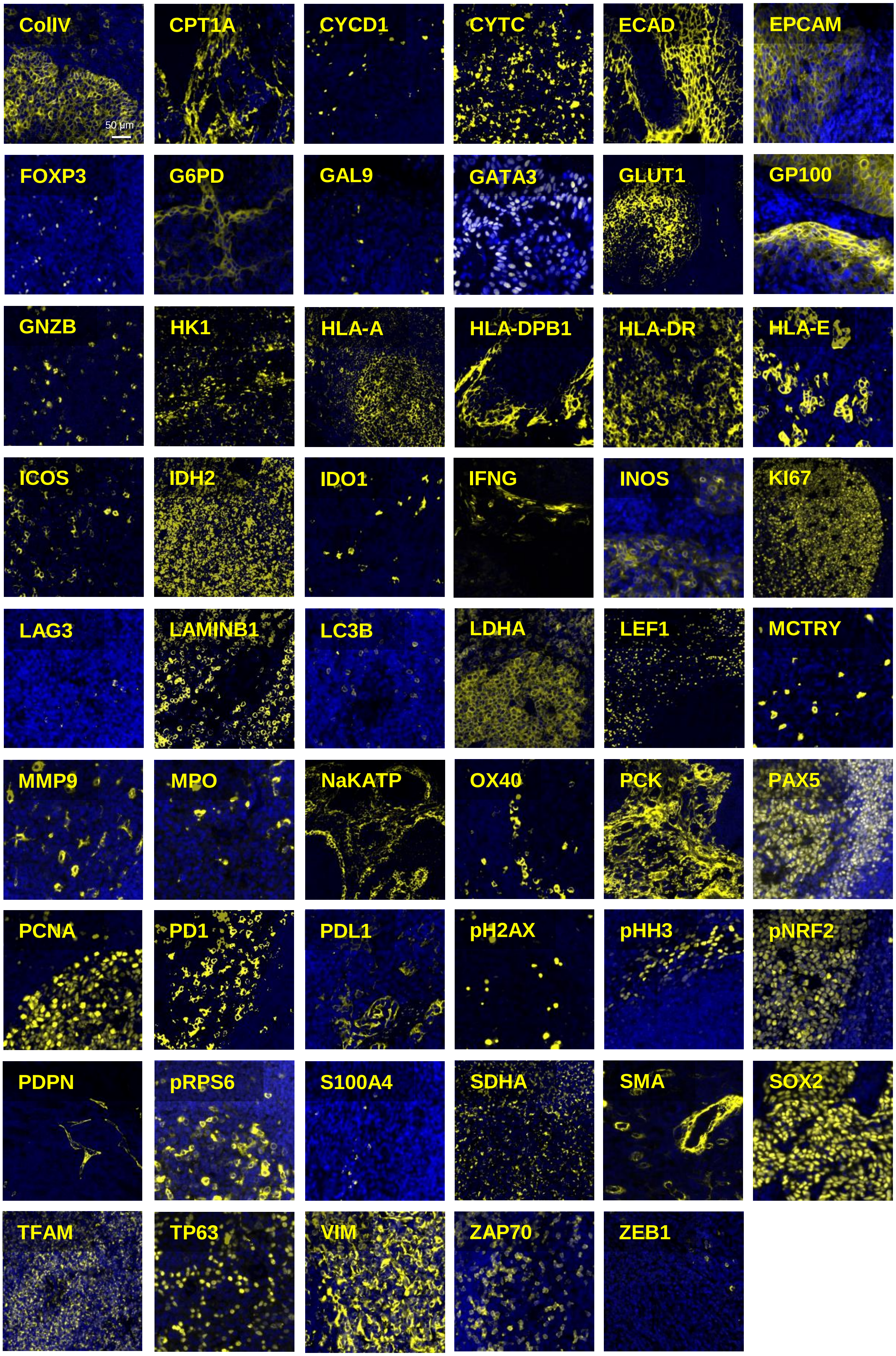


- Supplementary Figure 3 -

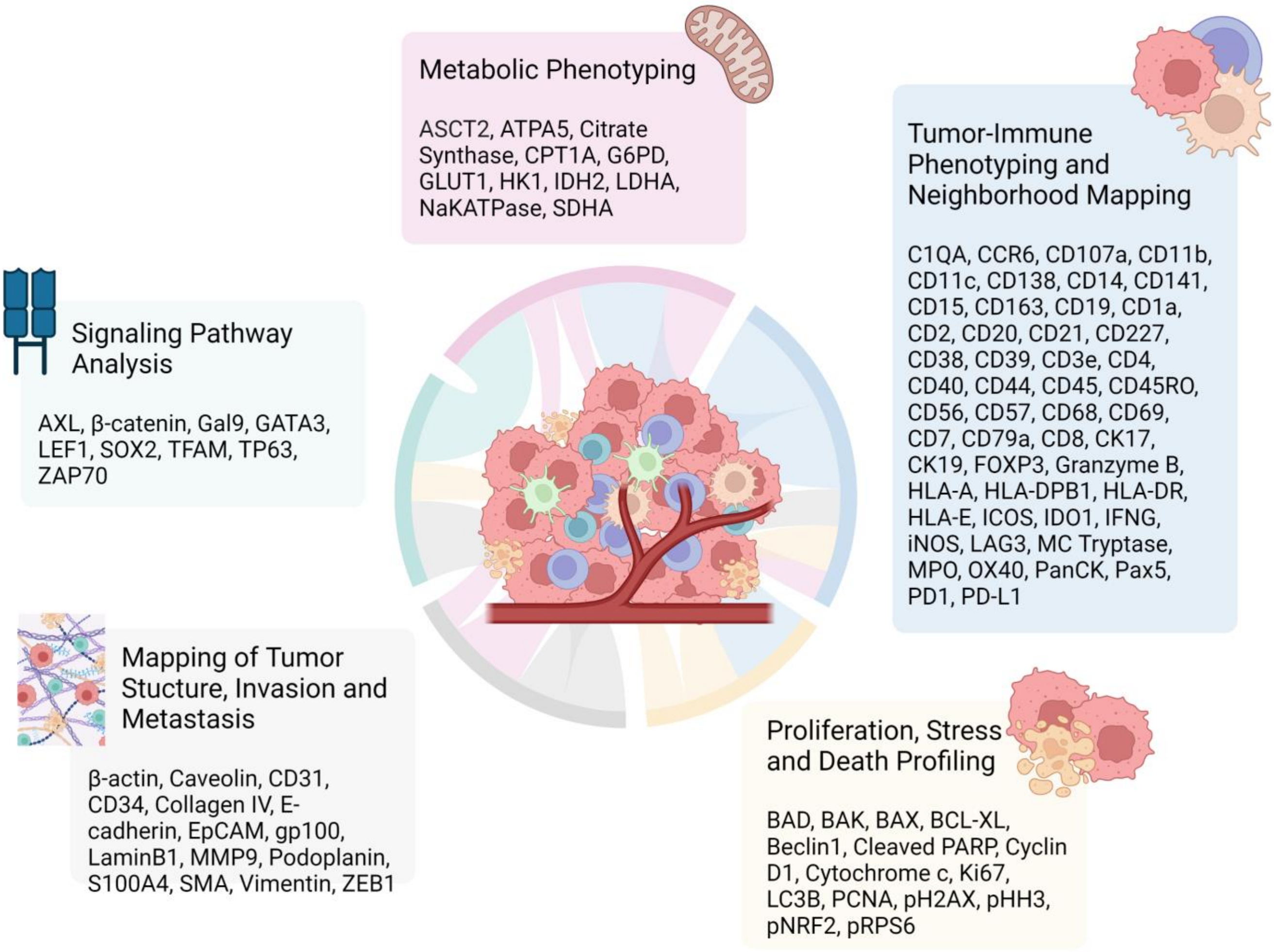


**- Supplementary Figure 4 -**

**A)** Metabolic and Stress Expression Profiles in Immune and Tumor cells

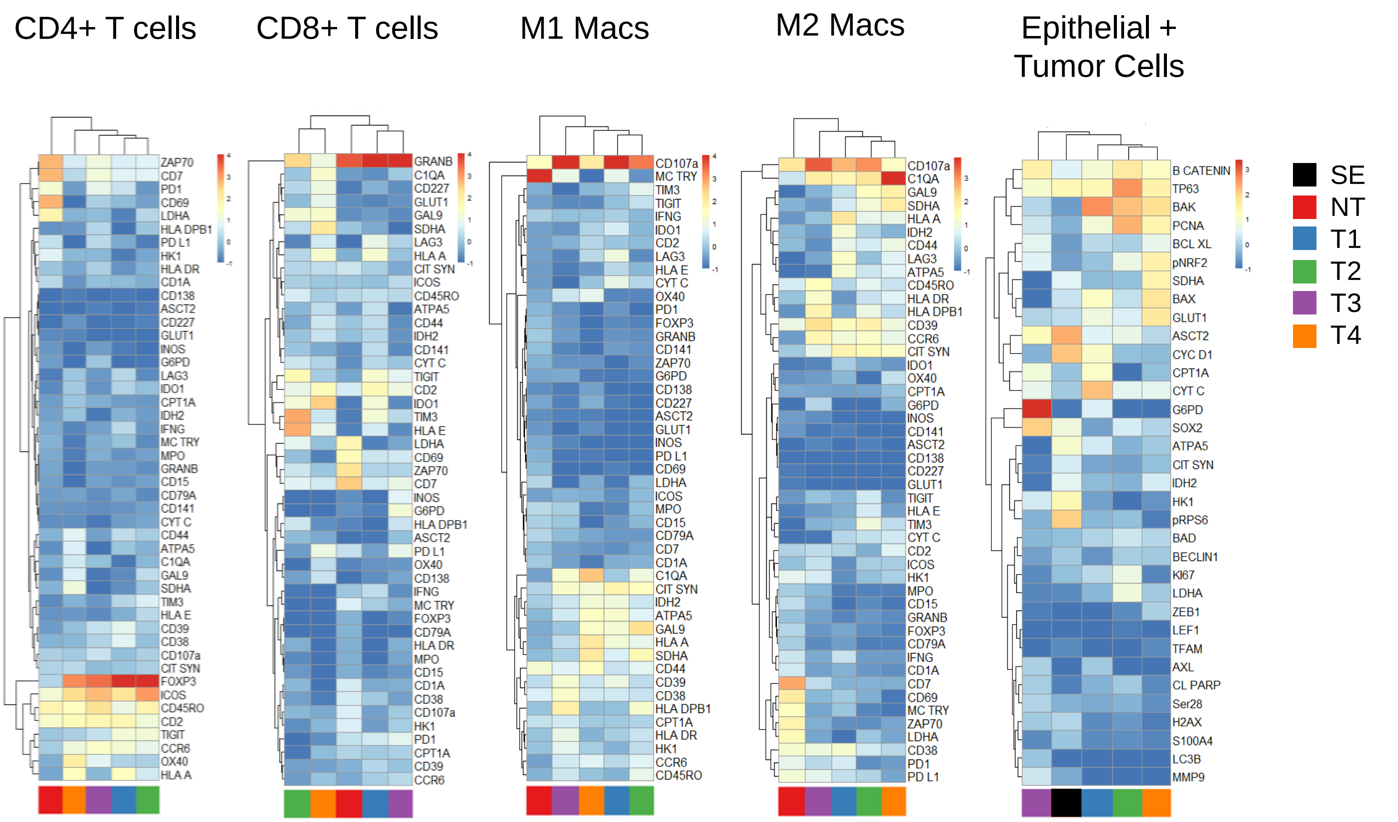


**B)** Cell-type Specific Expression of Metabolic and Stress Markers

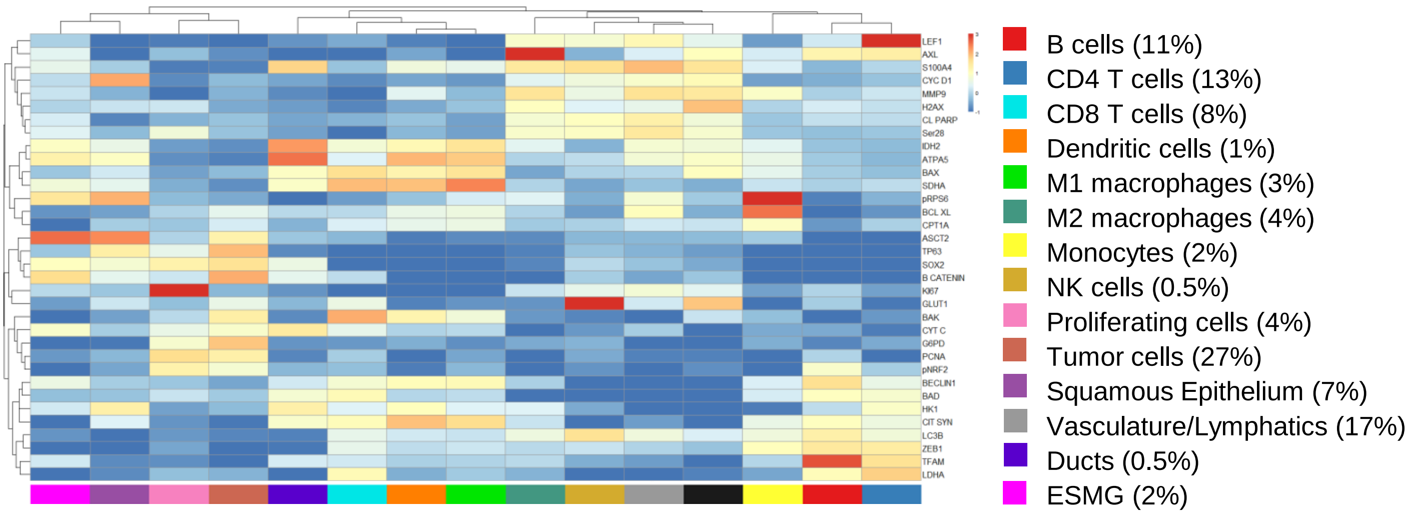


**- Supplementary Figure 5 -**
